# Simulated metabolic profiles reveal biases in pathway analysis methods

**DOI:** 10.1101/2025.03.27.645696

**Authors:** Juliette Cooke, Cecilia Wieder, Nathalie Poupin, Clément Frainay, Timothy Ebbels, Fabien Jourdan

**Author notes:** Contributing authors.

## Abstract

Initially developed for transcriptomics data, pathway analysis (PA) methods can introduce biases when applied to metabolomics data, especially if input parameters are not chosen with care. This is particularly true for exometabolomics data, where exported metabolites may be far from internal disruptions in the organism. Experimentally evaluating PA methods is practically impossible when the sample’s ”true” metabolic disruption is unknown. Using *in silico* metabolic modelling, we simulated metabolic profiles for entire pathway knockouts, providing both a known disruption site as well as a simulated metabolic profile for use in PA methods. PA should be able to detect the known disrupted pathway among the significantly enriched pathways for that profile. Through network-level statistics, visualisation, and graph-based metrics, we show that even when a given pathway is completely blocked, it may not be detected as significantly enriched with PA. This work highlights how some metabolomics data may not be suited to typical PA methods, and serves as a benchmark for analysing, improving and developing new PA tools.

## 1 Introduction

Pathway analysis (PA) methods aim to extract functional information from predefined sets (pathways) of molecular entities. In general, PA is based on calculating whether a given subset of differential molecules, usually profiled via high-throughput experiments, are significantly present among these existing groups. PA methods were initially developed for transcriptomics data (e.g. RNA-seq and microarrays) to uncover underlying patterns hidden in the large number of variables. Since then, PA methods have been applied to other research areas such as metabolomics (Chagoyen & Pazos, 2011; Karnovsky & Li, 2020; Marco-Ramell et al., 2018), where a list of perturbed metabolites (a metabolic profile) is functionally interpreted by identifying significantly enriched metabolic pathways. However, the widespread application of PA to metabolomics data can result in biases in interpretation, especially if input parameters are not chosen with care. Regardless of the origin of the data, the pathway database and the background set are important to define for PA, and can drastically affect the results from a single analysis. For metabolomics data, the biases in both compound detection and identification when using different analytical platforms can impact PA results in a way that is not currently taken into account in standard PA methods (Wieder et al., 2021).

Other biases can arise specifically with exometabolomics data, where the metabolic profile consists of extracellular metabolites measured in an external medium, such as cell culture media, or a biofluid like urine or blood. Although metabolomics and exometabolomics both use the same experimental approaches, they mainly differ in experimental design (e.g. sample setup and extraction) and interpretation. In exometabolomics, only metabolites that can be exported from the cell or produced externally can be measured in the extracellular medium. As a consequence, there can be many reaction steps between the measurable extracellular metabolites in the profile and internal disruptions in the system.

The underlying assumption in current applications of PA is that the metabolic profile measured in an exometabolome should reflect the pathways of the internal disrupted reactions. Biologically, there can be several biochemical steps (reactions) between the affected metabolic pathway and the metabolites detected in the exometabolomic profile, leading to potentially biased PA results. However, the community lacks ground truth datasets for benchmarking PA methods and validating the previous assumption. In the field of genomics, studies have defined hypothetical gold standard datasets, developed metrics for evaluating specific methods, and compared PA tools (Abatangelo et al., 2009; Liu, Dinu, Adewale, Potter, & Yasui, 2007; Nguyen, Shafi, Nguyen, & Draghici, 2019; Yu et al., 2017). For metabolomics, most PA evaluation approaches use existing experimental datasets with no known established ground-truth (Marco-Ramell et al., 2018; Wieder et al., 2021). To the best of our knowledge, no experimental or simulated datasets linking known metabolic disruptions and exometabolomics data exist.

For this purpose, we created a simulated dataset which associates known perturbations with simulated metabolic profiles. Using *in silico* metabolic modelling, we can simulate disruptions in genome-scale metabolic networks (GSMNs), creating a perturbed metabolic state. GSMNs are interconnected sets of metabolites, reactions, and metabolic genes, which together describe all known metabolic functions for a given organism. In particular, human GSMNs such as Recon2 (Thiele et al., 2013) and Human1 (Robinson et al., 2020) contain an existing set of pathway definitions for the reactions and metabolites from that network. In our dataset, we created the different known metabolic perturbations as entire (independently) knocked-out pathways. To be able to associate these perturbations with metabolic profiles, we used SAMBA (Cooke et al., 2024), a constraint-based modelling (CBM) method based on random flux sampling. By sampling the fluxes of exchange reactions (dedicated import and export reactions for each metabolite), SAMBA can be used to simulate the resulting metabolic profile for a given metabolic state, in this case a pathway knockout.

The dataset is therefore a compilation of known disruption sites (pathway knockouts), each associated with a simulated metabolic profile. The same hypothesis used in experimental PA can be used for our *in silico* dataset: PA methods should be able to detect the simulated known knocked-out pathway among the significantly enriched pathways for that profile (Fig. 1). The objective of this is to test the common assumption of PA: that exometabolomic profiles can be used to highlight an internal metabolic perturbation. In this paper, we use this benchmark dataset to assess to which extent PA methods are able to retrieve the correct perturbations, and to identify if network and pathway properties could help in identifying pathways which are more prone to misleading interpretations.

**Fig. 1:**
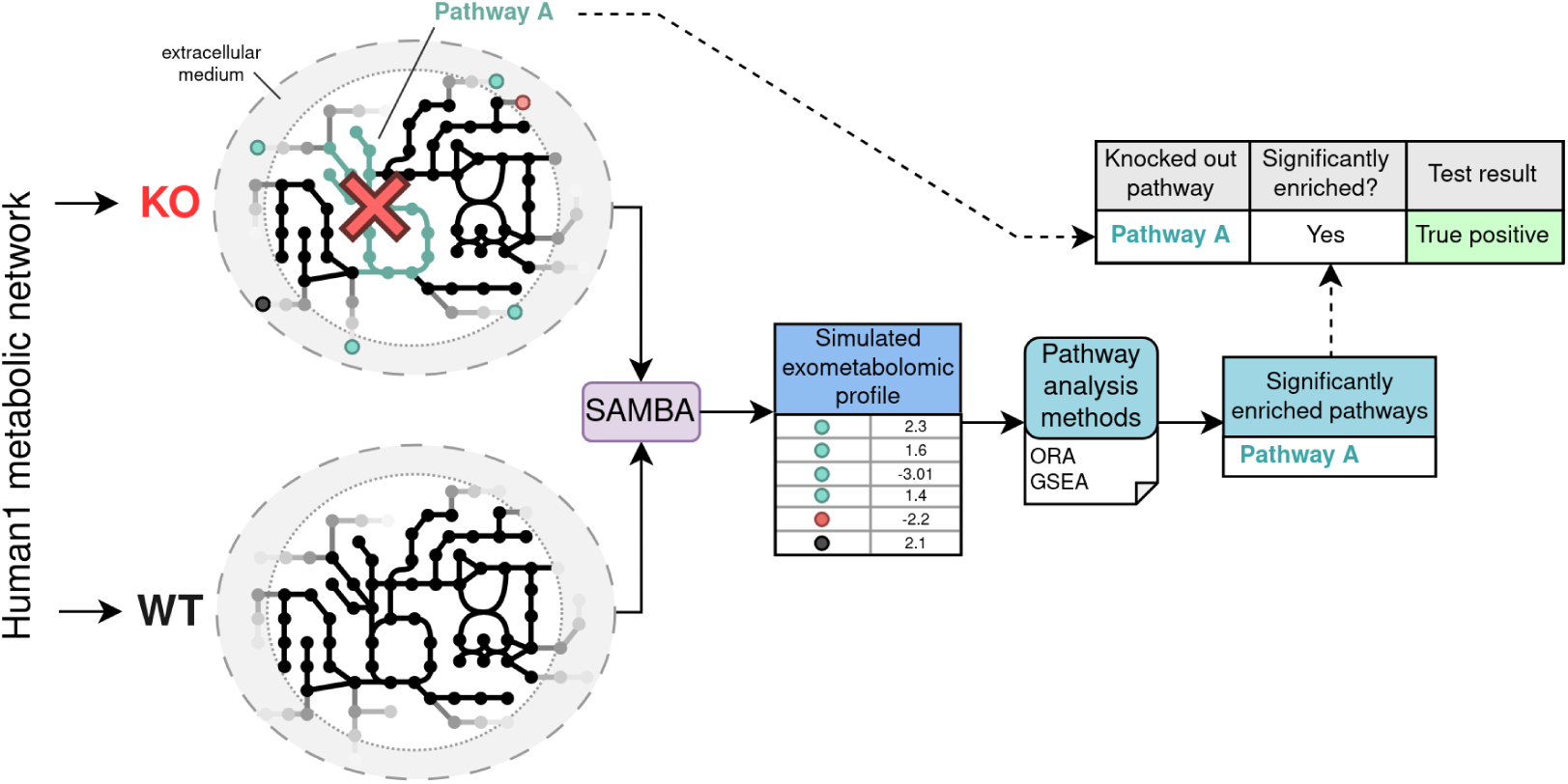
Graphical abstract of the creation of our pathway analysis benchmark dataset. Using the Human1 metabolic network, we knock out entire pathways, then simulate the corresponding exometabolomic profile. This profile can be used in pathway analysis methods, and the resulting significantly enriched pathways are then compared with the original pathway disruption.

## 2 Methods

### 2.1 Model and pathway definitions

In this work, we will present results obtained on Human1 (Robinson et al., 2020), the latest human GSMN. Human1 version 1.11 (https://github.com/SysBioChalmers/Human-GEM/releases/tag/v1.11.0) contains 142 metabolic pathways. Pathways in this model are defined on reactions, and a given reaction can only be present in one metabolic pathway. In a metabolic network, reactions have bounds: an upper and lower limit on flux, predefined in the model. We removed all blocked reactions in the models (reactions that carry no flux in the default model solution), and pruned the metabolites that were no longer tied to any reactions after this removal. This removed two pathways: Alkaloids biosynthesis and Triacylglycerol synthesis, which contained 2 and 1 reactions respectively, prior to pruning. We then converted the reaction pathway sets to metabolite pathway sets by extracting, for each reaction, its substrates and products and assigning them to that reaction’s pathway. Side compounds (such as H2O, ATP, NAD…) are included in the pathway sets, but were excluded for some analyses (metabolite overlap, graph metrics). Each metabolite can therefore be present in multiple pathways (see Fig. F in S1 Text for metabolite distributions across pathways). All main analyses were also run on another human metabolic network reconstruction, Recon2.2 (Swainston et al., 2016): the corresponding figures are shown in the Supplementary Material.

Pathways were then grouped into pathway super-categories by matching pathway names to KEGG (Kanehisa & Goto, 2000) pathways if possible, and using the KEGG pathway category names (https://www.genome.jp/kegg/pathway.html) (we were able to do this for 112 pathways). The 30 remaining pathways were manually matched to an existing category if possible, otherwise a new category was assigned to the pathway. The file containing all pathway categories, as well as all other data related to this work, is available at https://zenodo.org/records/14980037 and https://github.com/juliette-cooke/simulatedPA.

Among the pathway categories is the Miscellaneous category, containing pathways such as “Artifical reactions” and “Transport reactions”. For the purpose of this study, we excluded the 8 Miscellaneous pathways in Human1 as they were not deemed biologically relevant to knock out or analyse. They are mainly important for the network as a whole and for simulation purposes (e.g. producing metabolite pools, biomass, transporting metabolites across compartments), but not useful for PA since they are not actual metabolic pathways in the biological sense. It should also be noted that transport and exchange reactions in the network may be poorly curated, as there could be alternatives to well-known transporters (creating non-biological shortcuts), missing reactions, and incorrect assignments.

### 2.2 Simulation

To create the metabolic disruptions in the Human1 network, we knocked out entire pathways independently. Knocking out a pathway involves setting the bounds of every reaction in that pathway to [0,0]. Each pathway knockout is an independent model state and is simulated in a separate run, and is compared to the same WT (wild type) state, consisting of the same model in its default state for all runs.

To create metabolic profiles from pathway KOs, we applied SAMBA (Sampling Biomarker Analysis) method (Cooke et al., 2024). SAMBA uses these modified single pathway knockout models to simulate the fluxes in both the default WT state and all of the KO states using random sampling. SAMBA then compares the exchange reactions fluxes between the two states, and outputs a simulated metabolic profile based on the difference between the two states for each exchange reaction. Each exchange reaction in the network is assigned to a single metabolite, and its flux corresponds to the amount that the metabolite is being imported or exported. The increase or decrease of flux through an exchange reaction between KO and WT conditions translates to a change in the corresponding extracellular metabolite that can be measured. A z-score is calculated for each exchanged metabolite and describes both the direction and intensity of change between the two states. Therefore, SAMBA outputs a metabolic profile in the form of a list of z-scores for each pathway knockout state, which can be used in the same manner as an experimental metabolic profile in PA methods.

Some pathways, when knocked-out in the network, render the model infeasible, or unable to produce a minimum amount of biomass (set to 10% of the maximum biomass production for our simulations). The biomass reaction is shown in Eq A in S1 Text. The infeasible pathways are shown in Fig. 2 in red and in Fig. 3 as dotted light grey circles. The blocked pathways (no flux can be carried by them) are shown in grey in Fig. 2D. These 20 pathways (infeasible (18) and blocked (2)) were not able to be used in the subsequent analyses in this work.

**Fig. 2:**
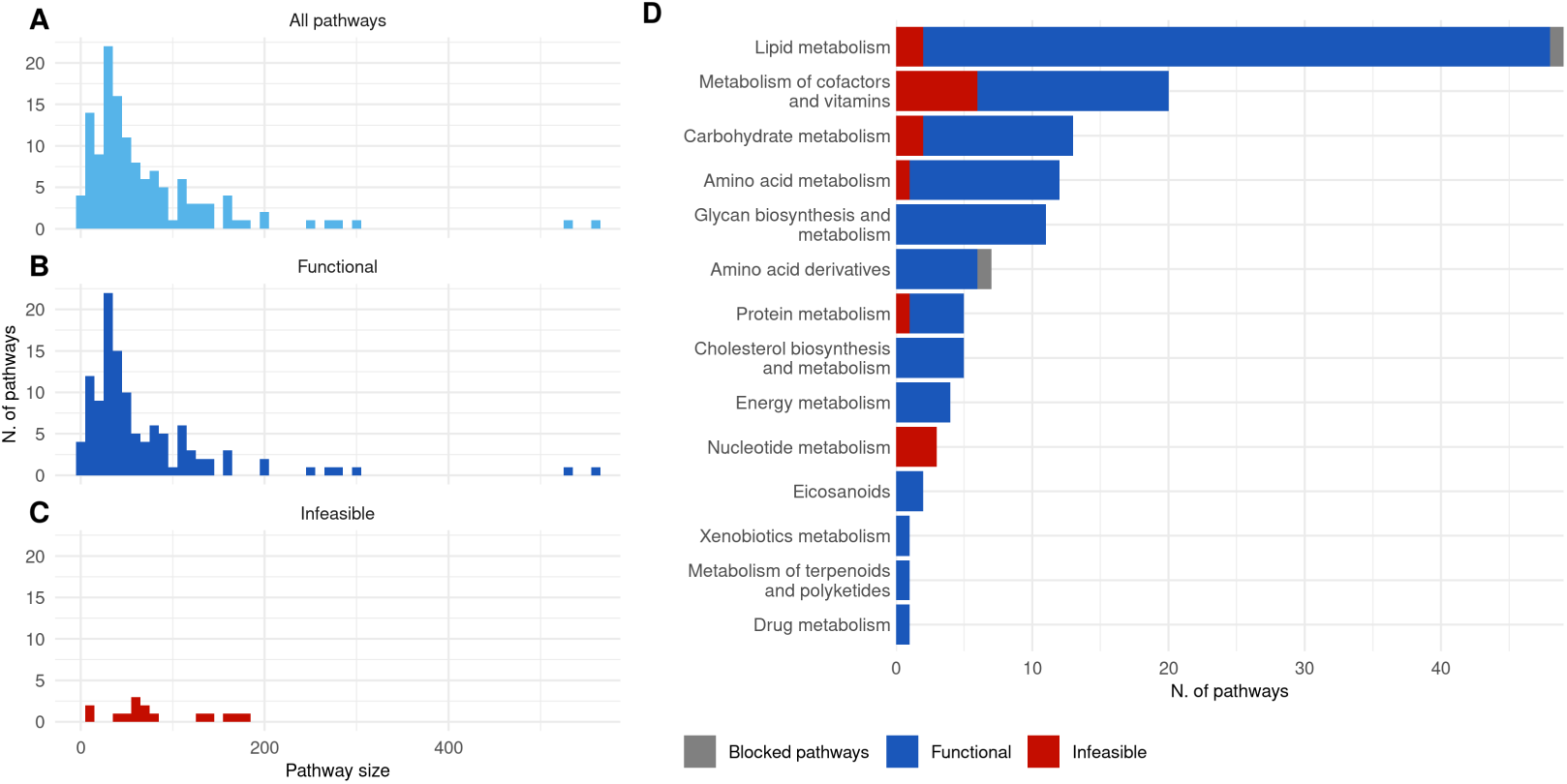
Pathway summary statistics. (**A**) Pathway sizes (number of metabolites) across all pathways in the Human 1 metabolic network, excluding miscellaneous pathways. (**B**) Sizes of pathways for which, when knocked-out, the model is able to produce biomass (functional). (**C**) Sizes of pathways for which, when knocked-out, the model is unable to produce biomass (infeasible). (**D**) Number of pathways per pathway category, coloured by whether the model is able to produce biomass or not when each pathway is knocked-out.

**Fig. 3:**
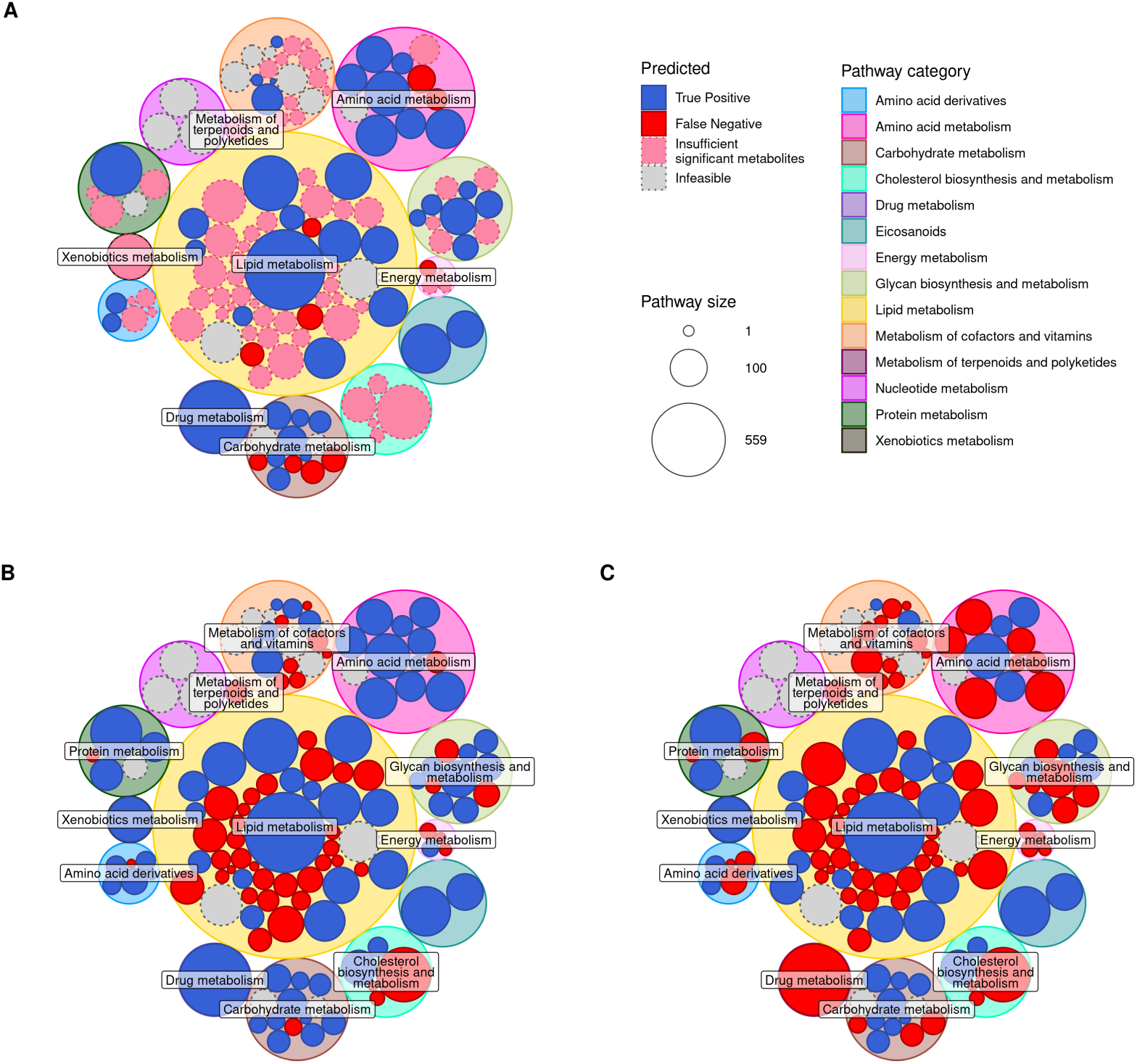
Circle pack plots for each pathway analysis method on the simulated dataset, (**A**) ORA, (**B**) GSEA with absolute (unsigned) z-score values as input, and (**C**) GSEA with raw (signed) z-score values as input. Major circles are coloured according to the pathway category. Minor circles are sized by pathway size, and coloured by significantly enriched when KO’d (TP), not significantly enriched when KO’d (FN), not run in ORA (due to insufficient significant metabolites), and infeasible when KO’d (as seen in Fig. 2).

### 2.3 Pathway analysis

Two PA methods were evaluated: over-representation analysis (ORA) and gene set enrichment analysis (GSEA) (Subramanian et al., 2005). ORA was implemented via the `ssPA` Python package (Wieder, Lai, & Ebbels, 2022) and GSEA was implemented via the `GSEApy` Python package (Fang, Liu, & Peltz, 2023).

Prior to ORA, the z-scores were converted to p-values using the survival function implemented by the `scipy` package (equivalent to 1 − *CDF* (cumulative distribution function)). The resulting p-values were then thresholded at *p* ≤ 0.05 to identify differential metabolites (no multiple testing correction, to preserve a minimal amount of significant metabolites). The background set was defined as all metabolites which could be exchanged (total 1,476, out of 4,156 unique metabolites in the model). These were then input to ORA for pathway analysis using the right-tailed Fisher’s exact test. The minimum threshold for a pathway to be tested was set to 3 significant metabolites per pathway.

GSEA was run using a pre-ranked list of metabolites, where either the signed or absolute z-score was used to represent each metabolite’s rank. GSEA was run with default parameters (1000 sample permutations). Significant pathways for both ORA and GSEA were defined as those with *p* ≤ 0.05 (no multiple testing correction, again, to preserve a minimum amount of signal).

### 2.4 Performance evaluation

To evaluate each PA method’s performance, we assigned the KO’d pathway tested in each KO scenario as either a true positive (TP) or false negative (FN). Therefore, each pathway can only be a TP or FN once across all scenarios: in the scenario where its reactions are knocked-out. Then, each additional pathway for each KO scenario was assigned either false positive (FP) or true negative (TN) depending on its enrichment significance. These were counted to form the TN and FP raw counts for each pathway knockout. They were defined as follows:

- TP: The target pathway containing the KO reactions and detected as enriched from the exometabolomic profile (*p* ≤ 0.05)
- FN: The target pathway containing the KO reactions but not detected as enriched (*p >* 0.05).
- FP: A pathway not containing the KO reactions and detected as enriched from the exometabolomic profile (*p* ≤ 0.05)
- TN: A pathway not containing the KO reactions, and not detected as enriched (*p >* 0.05)

The rates for each metric were then calculated across all KO scenarios in order to gain an average view of each method’s predictions. The TP and FN rates (TPR and FNR) across all KO scenarios are between 0 and 1 since they are calculated as averages of 0s and 1s for each pathway knockout. The numbers of FP and TN in each scenario can be between 0 and 117. We calculate their rates (FPR and TNR) as the average number of FP and TN pathways across all knockout scenarios. They are defined as follows, where *N* is the number of tested pathways (number of scenarios, *N* = 118 in this case), and *i* is a given scenario:

- TPR: the proportion of TPs across all scenarios (calculated as an average since *TP_i_* can only be 0 or 1):

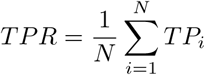

- FNR: the proportion of FNs across all scenarios:

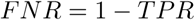

- FPR: average number of FPs across all scenarios:

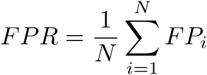

where *FP_i_*is the number of FP pathways in scenario i.

- TNR: the average number of pathways which do not fall into the other metrics:

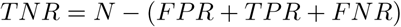

When calculating the FNR for ORA results, this method excludes pathways not testable (due to insufficient number of differential metabolites present to run the calculation (less than 3)). These non-testable pathways are then added back in, and the TPR and FNR are adjusted relative to the extra pathways. The non-testable pathways are shown as a new metric in a separate colour (pink) in the Results section plots.

### 2.5 Visualisation

All R plots were made using R 4.2.2, ggplot2, ggraph and other related packages, which can be found in the public notebooks at https://github.com/juliette-cooke/simulatedPA and https://zenodo.org/records/14980037.

The Met4J toolbox (https://forgemia.inra.fr/metexplore/met4j) was used to create the weighted pathway-level networks from SBMLs. The `PathwayNet` feature creates a graph where nodes are pathways and two nodes are conneced by an edge if the pathways share at least one metabolite, and sets as the edge attribute the number of shared metabolites between each pair of pathways, excluding side compounds. A heatmap showing the overlap between pathways is shown in Fig. E in S1 Text. The file containing the list of side compounds can be found in https://github.com/juliette-cooke/simulatedPA. The weights are based on the number of metabolites each pair of pathways shares. This weighted undirected graph was then imported into Cytoscape (Shannon et al., 2003), to both visualise the pathway graph and to calculate the graph-based statistics for each pathway.

### 2.6 Pathway metrics

The *exchangeable ratio* is defined for a pathway by the number of unique metabolites, not counting side compounds, in that pathway which are able to be exchanged (have an exchange reaction somewhere in the network) divided by the total number of unique metabolites in that pathway (again not counting side compounds). This penalises side compounds as even if they are exchangeable, they usually do not add much information to the metabolic profile for enrichment.

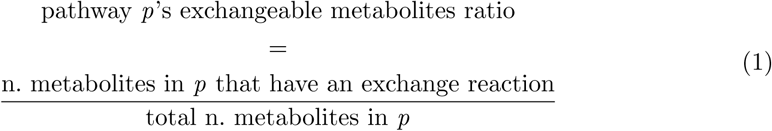

The metabolic profile *uniqueness score* is calculated for a given metabolic profile. A pathway’s *uniqueness score* is therefore the score associated with its metabolic profile when knocked out. This score includes in the metabolic profile only metabolites which pass a given z-score threshold (thr=2), and counts how many metabolites are in the resulting list across all pathway knockouts. Then, for each pathway KO, the counts are inversely summed, leading to a weighted score which rewards unique metabolites:

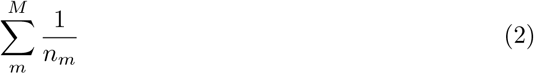

where *m* is each metabolite in that profile, *M* is the last metabolite, and *n_m_*is the number of times metabolite *m* appears across all metabolic profiles. The higher the *uniqueness score*, the more unique the metabolic profile is.

## 3 Results and Discussion

### 3.1 Human1 pathways summary statistics and pathway knockout feasibility

An initial analysis was performed to describe the pathway set extracted from the Human1 metabolic network. For each pathway, its size in number of metabolites is reported in Fig. 2A. When certain pathways are knocked out in the network, the model is unable to produce the minimum required amount of biomass (see Methods). These 18 infeasible pathways are shown in red in Fig. 2C and D.

Each pathway was sorted into a pathway super-category according to KEGG pathway groups (see Methods). The distribution of pathways in these categories is shown in Fig. 2D, along with the ratio of functional/infeasible when knocked out, as well as blocked pathways in grey. Certain categories contain many infeasible pathway knockouts such as Nucleotide metabolism, Protein metabolism, Metabolism of cofactors and vitamins, and Carbohydrate metabolism (Fig. 2), which seems relevant biologically as these categories are the sole requirements for producing several metabolites necessary for biomass production (see Fig. A in S1 Text). Lipid metabolism encompasses a large number of pathways, only a few of which are required to be active to produce biomass. The essential (infeasible) lipid pathways are responsible for producing some of the cofactor pool biomass and some of the lipid pool biomass.

### 3.2 Evaluation of pathway analysis methods on the simulated dataset

Classically, when using PA methods, the goal is to unveil the perturbed pathways in the studied condition. Here, each pathway which is able to be knocked out in the model constitutes a “truth” result that can be benchmarked with PA. A metabolic model containing a knocked-out pathway can then be used with SAMBA (Cooke et al., 2024) to simulate a metabolic profile associated with that knockout. For each functional (able to produce biomass when knocked out) pathway knockout, its corresponding simulated metabolic profile was used in ORA and GSEA as described in Pathway analysis (with two different inputs for GSEA: signed (raw) and unsigned (absolute) metabolic profiles). The significantly enriched pathways for each analysis were then evaluated as defined in Performance evaluation.

Fig. 3 shows the results of this evaluation for all pathways for each PA method. ORA requires a threshold on the input differentially abundant (DA) metabolites, resulting in 66 pathway knockout conditions for which there were not enough DA metabolites (pink in Fig. 3A) and therefore ORA was unable to be run (these could be considered FNs, but were kept separate for clarity). ORA has less FNs than the GSEA methods, and generally the same TPs as GSEA with absolute z-scores. The pathway knockouts with no metabolites which pass the DA threshold for ORA tend to also be FNs in GSEA. The ORA DA p-value thresholds can be seen in Fig. C in S1 Text.

When using signed z-scores as input for GSEA, predictions appear to be less precise than when using absolute (unsigned) z-scores in GSEA: many more pathways were unable to be detected as significantly enriched despite being knocked out in the model (Fig. 3B and C). The lipid metabolic pathways are the hardest to detect as significantly enriched across all three methods. These pathways appear to be some of the smaller lipid pathways, indicating a potential bias towards pathway size in PA methods.

The difference between the two GSEA approaches is intriguing. The hypothesis behind the original GSEA applied to transcriptomics is based on the assumption that genes are up- or down-regulated in sets in response to a disruption. Pathways are then defined based on these functional sets of genes. It is difficult to say whether we should apply the same assumption to metabolites, as a set of metabolites in a pathway may not all be over-abundant, but instead split across over- and under-abundance. Metabolic pathways are generally not defined on how correlated the behaviours of metabolites are, but tend to be defined around a specific function or metabolite. An example of this difference between signed and absolute z-scores using our simulated data for a specific pathway knockout is shown in Fig. B in S1 Text. Importantly, only the GSEA performed on absolute z-scores rather than signed z-scores was able to produce a significant p-value for some pathways. In the unsigned GSEA, conflicting z-score signs, reflecting metabolite abundance, resulted in a lower enrichment score test statistic and hence insignificant p-values.

As a summary analysis of these results, Fig. 4A shows the proportions of correct enrichments (TP) and missed enrichments (FN) for each method. Fig. 4B shows the average number of pathways that were either significantly enriched (FP) or not (TN), despite not being knocked out in that corresponding metabolic state. Note that each KO’d pathway can become either a TP or FN, whereas TN & FP rates are computed as the mean proportions over the 118 knock outs (see Performance evaluation).

**Fig. 4:**
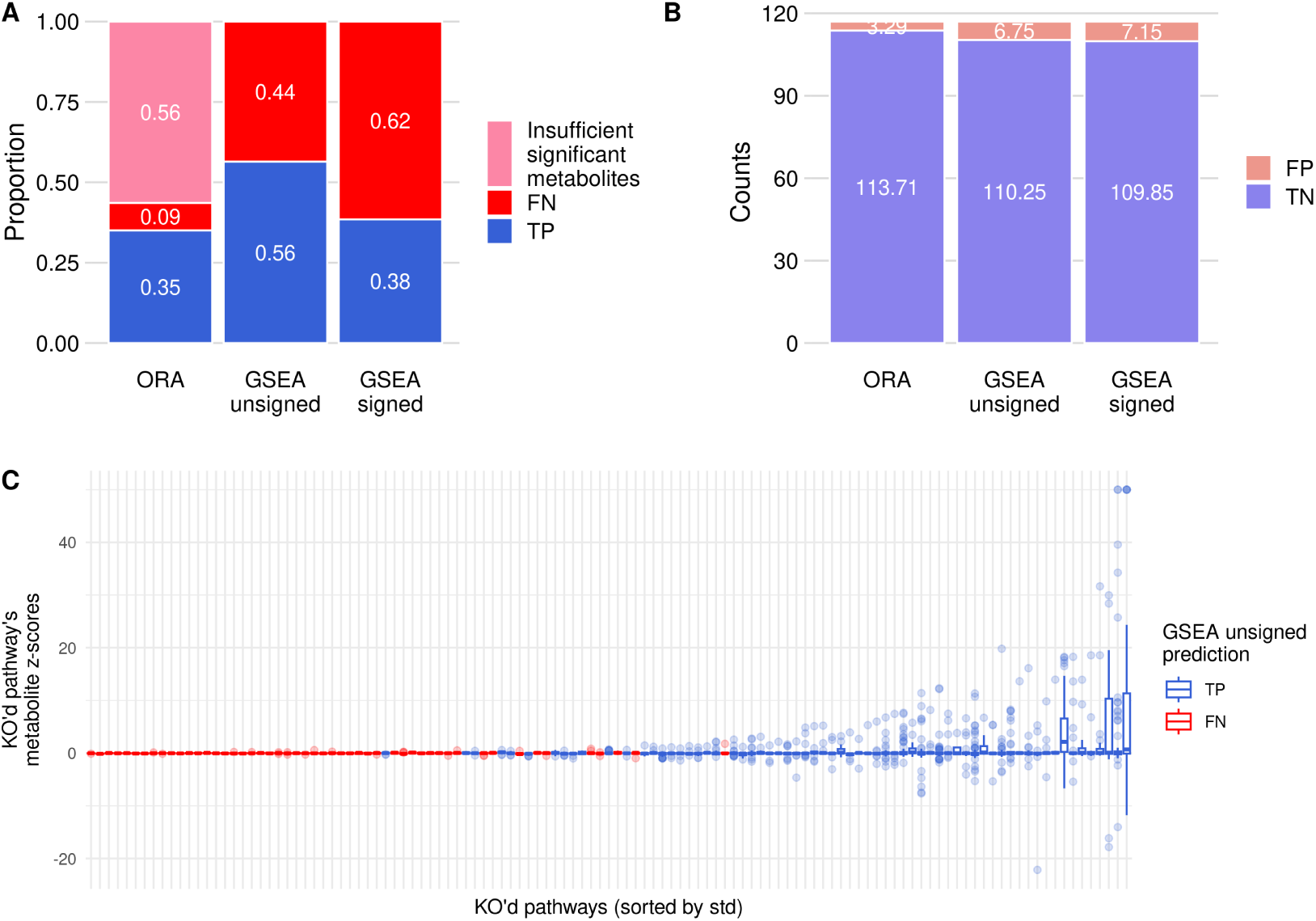
(A) Pathway enrichment summary proportions, for each tested PA method. TP pathways are shown in blue, FN pathways are shown in red, and pathway KOs for which there were no DA metabolites are shown in pink. (B) Pathway enrichment summary counts, for each tested PA method. TN pathways are shown in light blue, and FP pathways are shown in light red. (C) Boxplots of metabolite z-scores for each knocked out pathway. Pathways (x-axis) are sorted by standard deviation, and coloured by GSEA prediction (TP/FN). The y-axis shows the z-scores of all metabolites in that knocked out pathway. Outliers are shown as points.

The percentage of TPs in ORA is 35%, meaning that 65% of knocked out pathways were not significantly enriched due to either an insufficient number of DA metabolites (56%) or a non-significant enrichment (9%). Out of the ORA tests that were able to run (TP+FN), 80% were TP, which shows that due to the stringent filter on DA metabolites, those that do pass the filter tend to reflect the knocked out pathway’s disruptions correctly.

The percentage of TPs is 56% when using the unsigned metabolite z-scores in GSEA, and drops to 38% with the signed values in GSEA. Additionally, the false positive and true negative rates (Fig. 4B) are low and high respectively. This shows that for any given disruption, many pathways are not significantly disrupted and therefore enriched (TN), and there are around 3-7 pathways on average that were disrupted but were not the original knockout (FP). True negative pathways are pathways that were not knocked out in the simulation and for which there were not enough significantly disrupted metabolites for it to be significantly enriched. Here, false positive pathways are pathways that were sufficiently disrupted by the original pathway knockout to be significantly enriched, which can highlight the consequences of a knockout.

As a first step to understanding why some pathways are poorly detected as enriched, Fig. 4C shows the distributions of metabolite z-scores for each knocked out pathway. Pathways that are enriched when they are knocked out (TP, blue boxplots) tend to have a larger variation in metabolite z-scores (higher standard deviation), revealing a larger effect on their own metabolites. This is emphasized by a small number of extremely affected metabolites per pathway knockout, shown as outliers in the plot. Conversely, FN pathways (red boxplots) tend to have very little effect on the export and import of their own metabolites. Knocking out these pathways does not cause a large enough effect on their own metabolites to be significantly enriched when using the corresponding metabolic profile in PA methods.

### 3.3 Pathways are unequally detected as enriched when using the simulated exometabolomic profile

As shown in Figs. 3 and 4, some pathways, when knocked out, have a significant effect on their own metabolites, resulting in a TP enrichment. Others, despite also being knocked out, do not reach this level of metabolite perturbations, leading to a FN enrichment (in our case, these tend to be small lipid pathways). Generally, these FNs appear to be caused by a lack of effect on the specific metabolites belonging to the knocked out pathway (see Fig. 4C). Even those with the smallest effect across all metabolites have some variation in metabolite exchanges when knocked out (Fig. D in S1 Text), leading to at least one false positive.

This result in itself shows an inequality of each pathway with regards to PA methods, which test each pathway in a univariate manner. The FN pathways are interesting to analyse as they reveal potential biases in PA methods, which could cause erroneous interpretation in a biological experiment, as these are perturbations which may not cause enough metabolic disruptions in its own pathway to be detected via PA, but could still have an effect on metabolism.

Intuitively, when knocking out a pathway, other pathways are also affected, which in turn affects the metabolic profile for that condition. These additional significantly enriched pathways were not the original perturbation (FPs), but can still be interesting for biological interpretation. The FPs reveal that when several pathways are significantly enriched in a dataset’s PA results, some pathways may not be the origin of the perturbation, and may be side effects, which can both be intriguing but also misleading.

### 3.4 Intrinsic pathway properties as indicators of enrichment detectability

Following the evaluation of PA methods using our simulated dataset, we aimed to understand why certain pathways were not significantly enriched despite being knocked out in the network. To do this, we calculated various metrics and properties to understand both how pathways are defined as well as the relationships between them.

The exchangeable metabolite ratio was calculated for each pathway as described in Methods. It represents the proportion of metabolites in a pathway which can be transferred to the exometabolome through exchange reactions, and therefore how much a pathway can be represented in an exometabolomic profile. Fig. 5 shows this ratio for each pathway (blue bars), with its corresponding pathway category and the PA predictions (GSEA unsigned). A low exchangeable metabolite ratio implies that the pathway is poorly represented in terms of exported metabolites. This can be seen in Fig. 5 where the pathways with a higher exchangeable ratio tend to be TP, and those with a lower exchangeable ratio tend to be FN when using GSEA unsigned. This is a general trend with some exceptions, as there are some highly exchangeable pathways which were not enriched when knocked out (e.g. Heme degradation, Hippurate metabolism), and inversely, other pathways which were correctly enriched but with a low exchangeable ratio (e.g. Glycerolipid metabolism, Carnitine shuttle). In general, many of the Lipid metabolism pathways have a low exchangeable ratio, and “Glycan biosynthesis and metabolism” pathways are mostly below the halfway mark. Using this exchangeable ratio score shows that having less exported metabolites makes it harder to for a pathway to be significantly enriched, since the pathway sets are defined based on all metabolites (not just exchange metabolites).

**Fig. 5:**
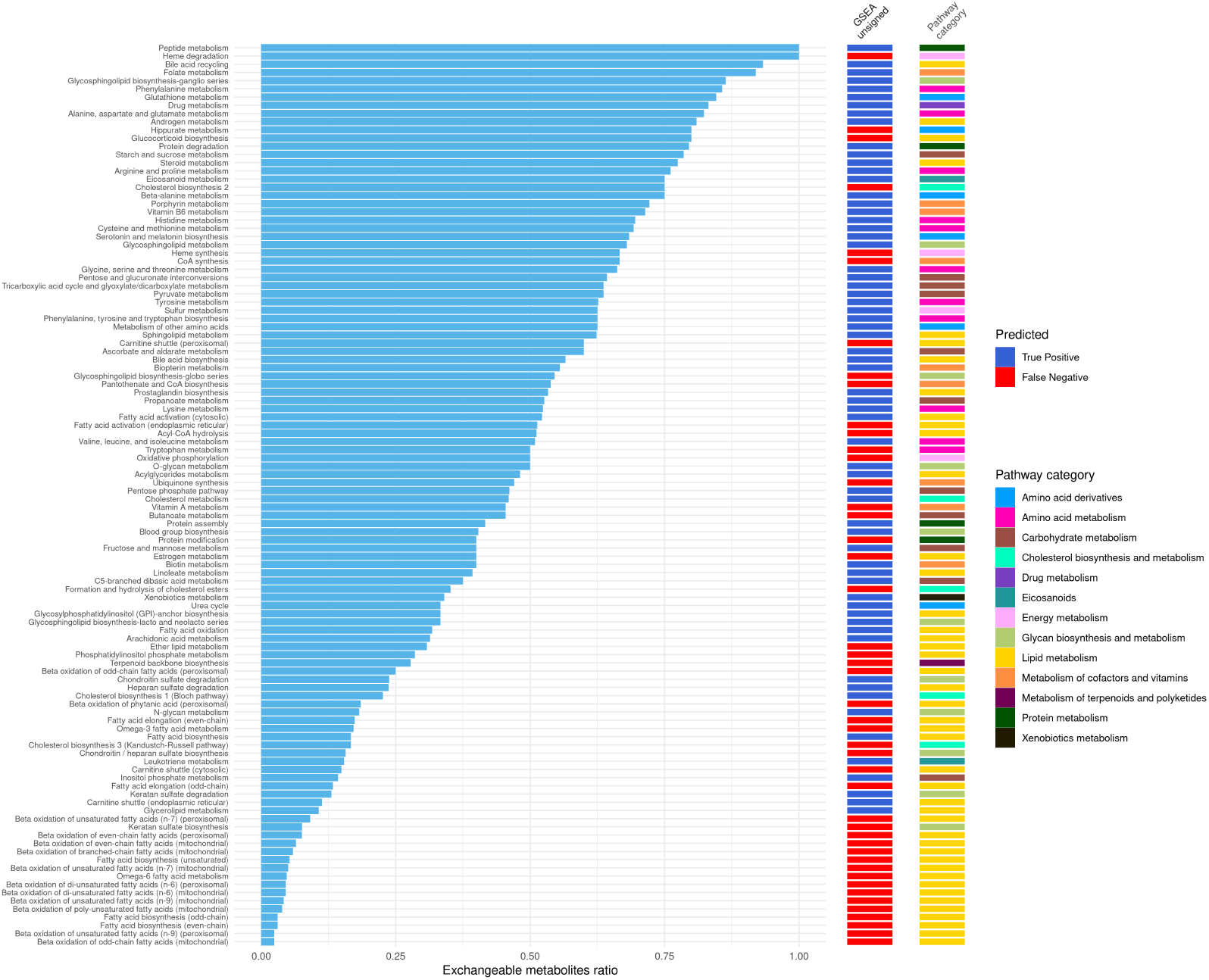
Exchangeable metabolite ratio for each tested pathway. The first column is the pathway’s category, and the next three columns are the enrichment results for each PA method.

A second score was calculated based on how unique each resulting metabolic profile is after knocking out each pathway (see Methods). A knocked out pathway’s corresponding simulate profile will have a high uniqueness score if many of its metabolites are not found in other pathway knockout profiles. Fig. 6A shows six intrinsic pathway metrics, for all pathways, separated into the two possible PA outcomes (TP and FN), for each PA method. The three metrics, pathway size, exchangeable ratio, and profile uniqueness score, are generally lower for FN pathways across methods, and GSEA unsigned pathway predictions are more separated compared to ORA and GSEA signed. We could assume that the smaller a pathway is, the smaller the impact when it is knocked out, since there are less total reactions being knocked out. The pathway size barplots in Fig. 6A show that while a pathway of any size can be a TP, those that are not enriched (FN) tend to be smaller, have lower exchangeable metabolite ratios, and lower uniqueness scores.

**Fig. 6:**
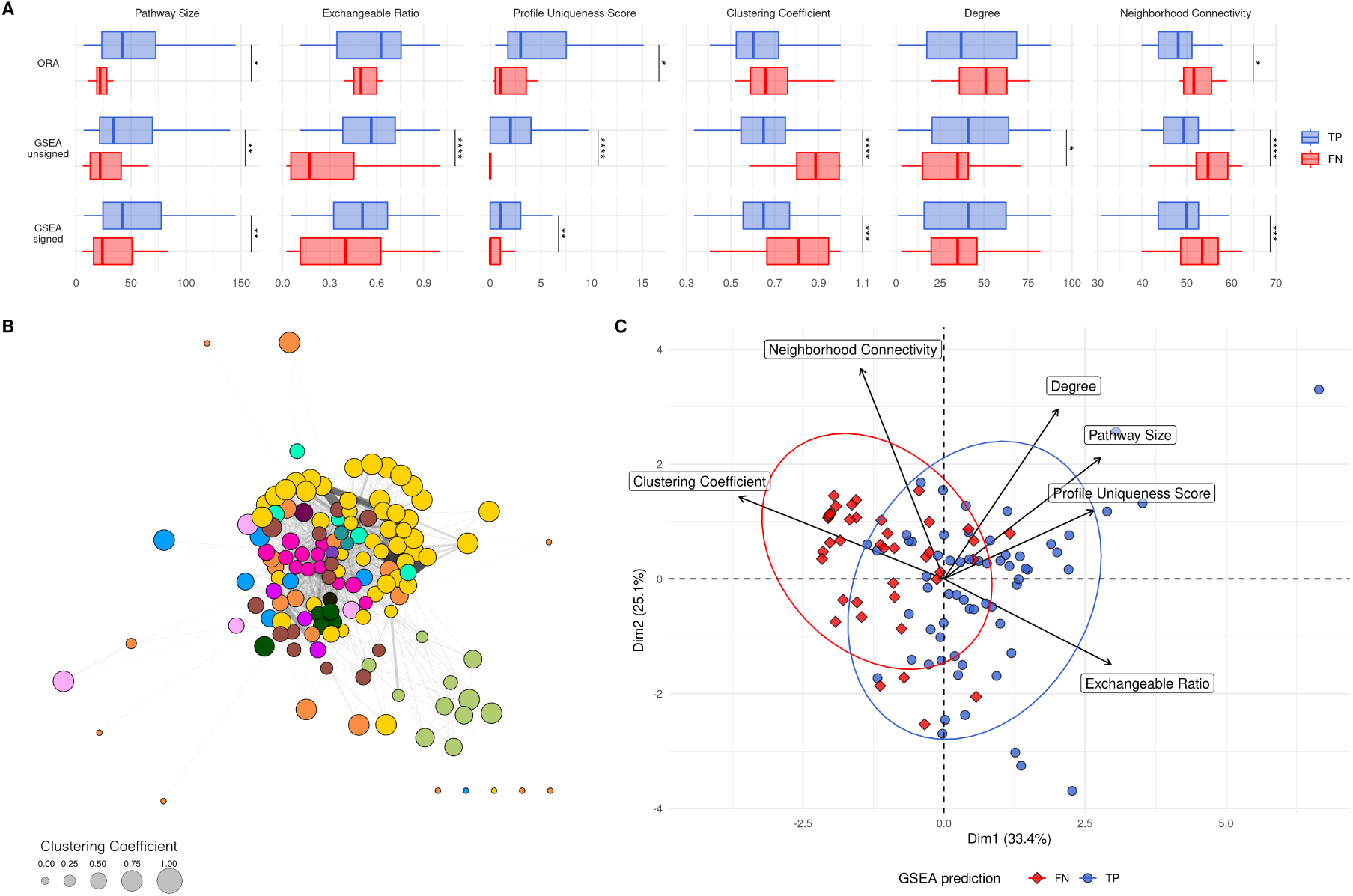
(A) Boxplots of pathway size, pathway exchangeable metabolite ratios, and pathway KO metabolic profile uniqueness score, as well as clustering coefficient, node degree, and neighbourhood connectivity. For each method, both the TP (blue) and FN (red) pathways have been separated. Wilcox tests (FDR adjusted) *: p ≤ 0.05, **: p ≤ 0.01, ***: p ≤ 0.001, ****: p ≤ 0.0001 (B) Pathway-level graph of all pathways in the Human1 metabolic network. Each node is a pathway, each edge is one or more shared metabolites between each pair of pathways. Node colour is the pathway category (see Fig. 5 for the legend), and node size is the clustering coefficient. (C) PCA of pathways using 6 metrics and properties. TP pathways are shown in blue (circles), FN pathways are shown in red (diamonds).

In order to gain more insight on how the metabolic network is structured into pathways, we computed the pathway-level weighted graph from the network (see Methods). A pair of pathways is connected if they share at least a metabolite, and the more metabolites they share (excluding side compounds), the higher the weight on the connecting edge. Fig. 6B shows this pathway graph, where each node is a pathway, coloured by pathway category. Using the edge-weighted spring embedded layout, pathways with a greater amount of shared metabolites are pulled together. The graph layout groups many of the pathways into their parent categories, such as Lipid metabolism, Glycan biosynthesis and metabolism, Amino acid metabolism, and Protein metabolism. The size of each circle reflects that pathway’s clustering coefficient, one of the graph metrics calculated from this graph. The clustering coefficient is shown along with the node degree and neighbourhood connectivity in Fig. 6A (see Methods in S1 Text for more details on these properties).

Together, these metrics and properties were assigned to each pathway for input into a PCA, shown in Fig. 6C. The combination of these properties tends to separate the TP pathways from the FN pathways, with most of the FN pathways being on the left half of the first principal component. FN pathways tend to have a higher clustering coefficient, a lower metabolic profile uniqueness score, and a lower exchangeable ratio (Fig. 6C). They also tend to be on the smaller side, although this is not a decisive factor, and are mainly lipid pathways (see Fig. G in S1 Text).

Since this approach depends on the calculated pathway properties, these conclusions will vary depending on the pathway database. To evaluate this effect, we ran all analyses on a second metabolic network, Recon2.2, as shown in Figs. H in S1 Text to K in S1 Text. These results show that the pathway properties we identified for Human1 do not discriminate as well as for Recon2.2 (especially pathway size), but still show a trend where FN pathways tend to have a lower exchangeable ratio for example. This emphasizes the fact that care must be taken when choosing the pathway database as basic properties can vary between them, despite PA methods treating them the same. Other pathway databases such as KEGG were considered through mapping Human1 IDs to KEGG compound IDs and using the KEGG pathway definitions. Figs. L in S1 Text and M in S1 Text show the same approach using these KEGG pathway definitions instead of those from Human1, all while using the Human1 simulated profiles. The proportions of TP/FN are similar to those with Human1 pathways, but these results should be analysed with caution. During the ID mapping, many metabolites were lost (only 506 of the 1559 exchange metabolites had KEGG IDs), and only 41 of the 142 Human1 pathways could be matched by name. This of course reduced the pool of pathways to analyse, but also introduced a bias where only the well-annotated metabolites and pathways were able to be taken into account. Since KEGG does not annotate metabolites explicitly as to exchangeability, as well as other cellular compartments, the rest of the analysis could not be carried out. We could therefore not confidently include it as a pathway source in our work. The KEGG network is also unable to be run as a functional metabolic model so could not be used for a standalone KO analysis. Future work could be focused on mapping more KEGG pathways (through reaction or gene IDs) and metabolites (through other IDs or manual curation) to Human1, which would then allow for a KEGG-based pathway KO simulation of metabolic profiles using Human1.

The trend of these metrics on TP and FN pathways is consistent with how current PA methods approach enrichment. A knocked out pathway is more likely to be enriched if its corresponding metabolic profile is more specific to that pathway, and if it contains metabolites that can be exported. Graph-based metrics are able to highlight the inherent links between groups of pathways which cannot be taken into account with current non topology-based PA methods. Current PA methods struggle to detect the correctly enriched pathway when knocking out this pathway has little effect on metabolite exports. In particular, ORA, due to its stringent threshold, misses many TPs but performs better on those that do pass the threshold. GSEA has a better proportion of TPs compared to ORA when taking all pathways into account, but only when using unsigned z-scores, and detects more FPs than ORA. Overall, the difference between the two GSEA methods is surprising, and could be examined further by future studies. The effect of the strict ORA threshold was expected, as ORA uses less of the metabolic profile than GSEA, and quantifying the difference between them is of value to the community.

Finally, the pathway metrics could be used to calculate an *a priori* confidence score for each pathway (or even areas in the network) before enrichment, describing how likely a given perturbation in that pathway is to lead to correct enrichment. Future work could provide these scores for pathways and pathway databases, which would help in choosing an appropriate database for PA, and could increase awareness of the biases these methods present, even leading to the development of more appropriate PA methods for exometabolomics data.

## 4 Conclusion

In this work, we show how effectively current pathway analysis methods can identify as enriched the pathway that is known to be disrupted, which is essential for communicating the risks of using pathway analysis for exometabolomics data. Care must be taken when interpreting pathway analysis results on exometabolomics data, especially when assuming the significantly enriched pathways are the origin points for the disruption, as some could simply be side effects. The simulated dataset used for this consists of metabolic profiles simulated for known pathway disruptions, and is a novel and direct link between what is expected to be enriched and what is actually enriched. The dataset we generated can also be used by the community to evaluate other existing PA methods, test newly created methods, and develop new ones more suited to metabolomics data. Our analysis of this work could also be the foundation for the development of a pathway-level score, indicating the level of confidence after detecting it as significantly enriched. Consequently, this work highlights how the nature of some metabolomics data may not be suited to typical PA methods, provides information on how difficult it may be for each pathway to be enriched depending on some inherent properties, and also serves as a benchmark for analysing, improving and potentially developing new PA tools.

## Acknowledgements

The authors thank Jake Bundy for his valuable discussions and input during the creation of this paper and the project. The authors are grateful to the GenoToul bioinformatics platform (Toulouse, France) for hosting and providing their computer cluster, used for running all sampling runs.

## Software and data availability

Data and code links:

- Github: https://github.com/juliette-cooke/simulatedPA
- Zenodo:

– Code: https://zenodo.org/records/14980037
– Data: https://zenodo.org/records/13753914

## Supplementary information

### S1 Text

#### Methods in S1 Text

The *clustering coefficient* quantifies how connected each pathway’s neighbouring pathways are. For a node, it is defined as the actual number of links between its neighbouring nodes divided by the maximum possible number of links that could exist between these nodes.

For a pathway *p_i_*:

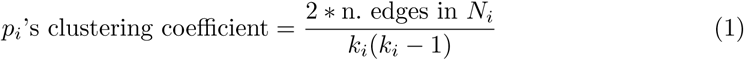

where *N_i_* is *p_i_*’s (immediate) neighbourhood, and *k_i_* is the number of nodes in *N_i_*.

The *neighbourhood connectivity* is defined as the average connectivity of all of a node’s neighbours. A node’s connectivity, or degree, is the number of connected edges, in our case the number of neighbours.

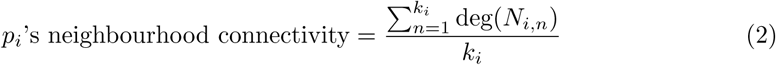

where *N_i_* is *p_i_*’s (immediate) neighbourhood, *N_i,n_* is the *n*th node in *N_i_*, and *k_i_* is the number of nodes in *N_i_*.

#### Eq A in S1 Text

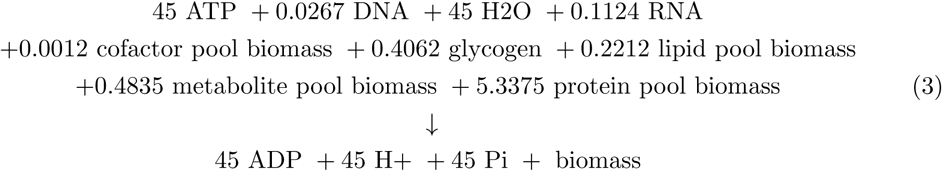

**Fig A in S1 Text:**
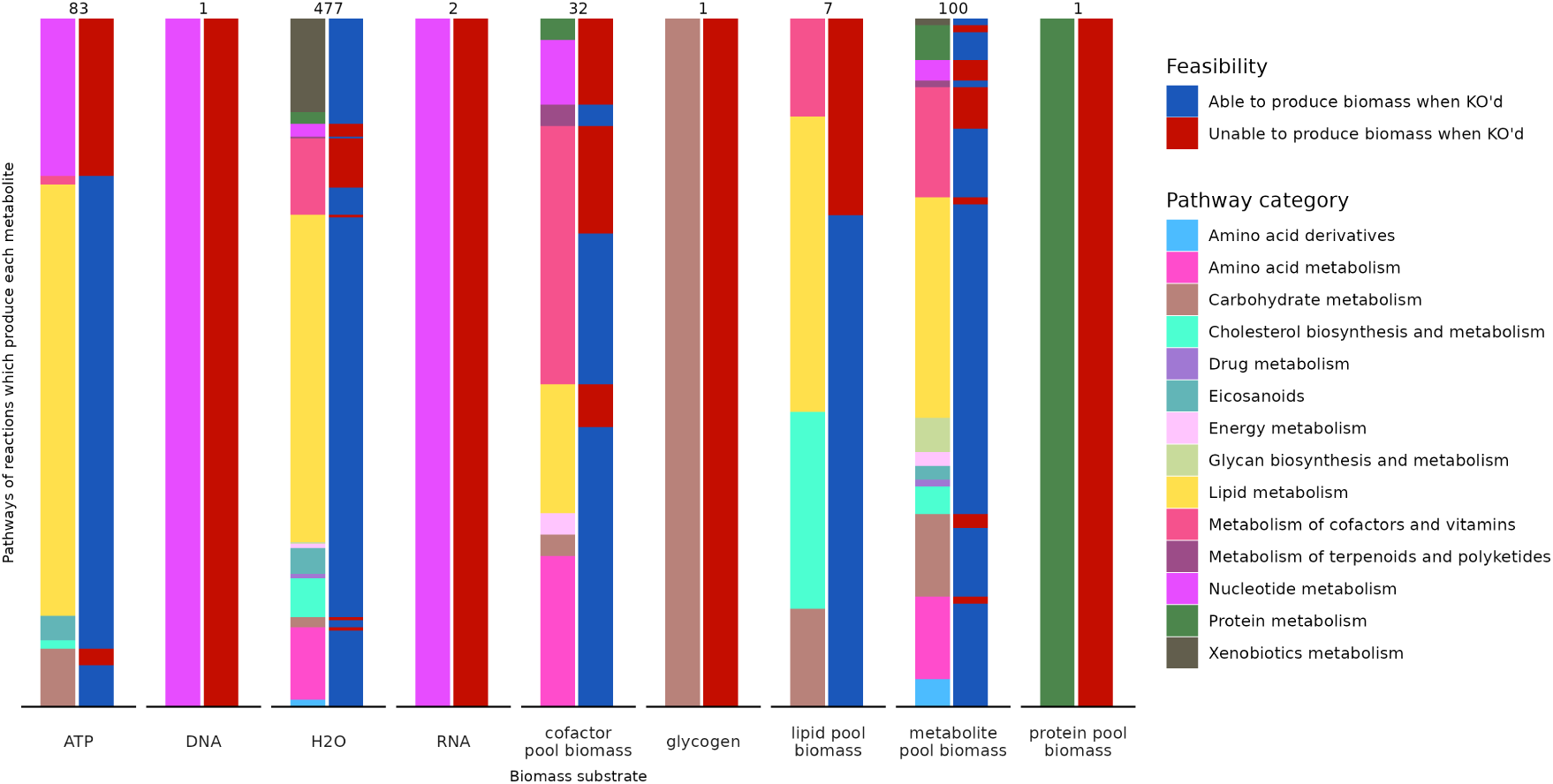
Pathway categories of reactions which produce each substrate in the biomass reaction. Each bar is the total number of reactions that produce that metabolite prior to the bioomass reaction. The left bars are the pathway super-categories for those reactions, and the right bars are whether those pathways are infeasible (red) or functional (blue).

**Fig B in S1 Text:**
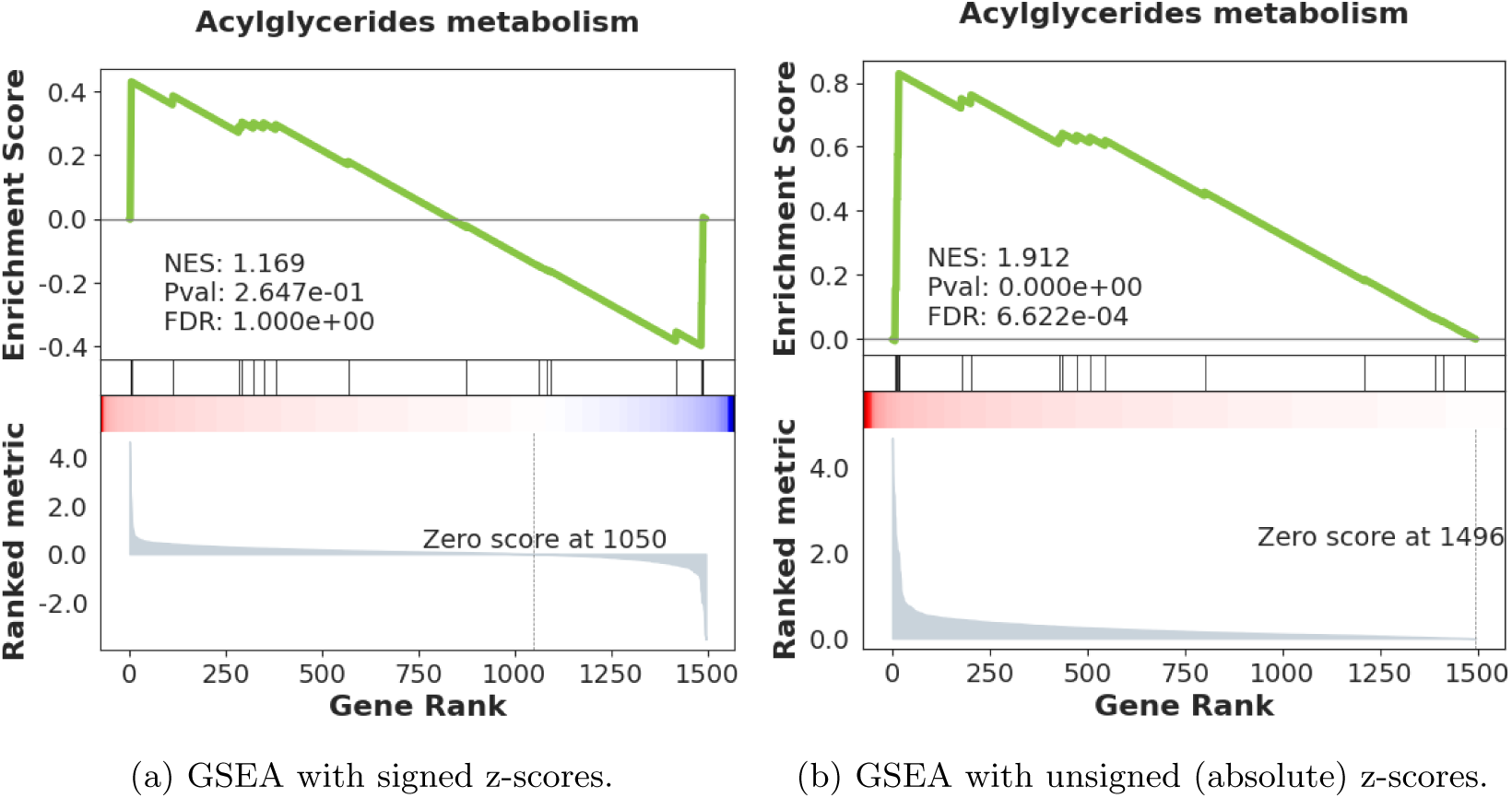
GSEA enrichment plots for Acylglycerides metabolism with GSEA abs and GSEA raw. The signed z-scores result in a split profile (similar contributions of high and low abundance metabolites), and a non-significant enrichment. By converting the z-scores to absolute values, they are clustered together in the ranking list, resulting in a correct significant enrichment.

**Fig C in S1 Text:**
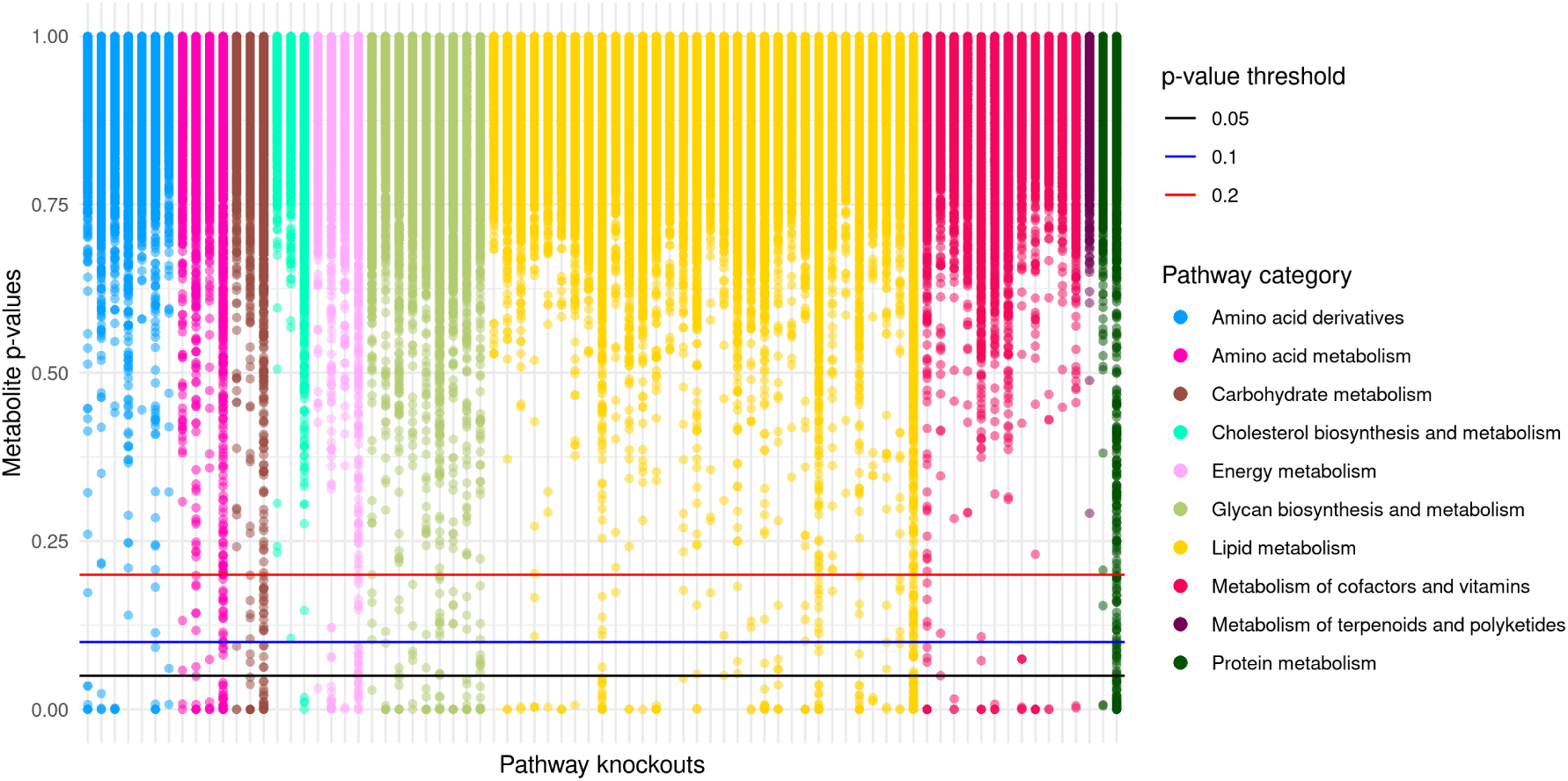
Different DA (differentially abundant) thresholds on all pathway z-scores for ORA. We used *p* ≤ 0.05 in our analyses. Pathways are grouped by super category.

**Fig D in S1 Text:**
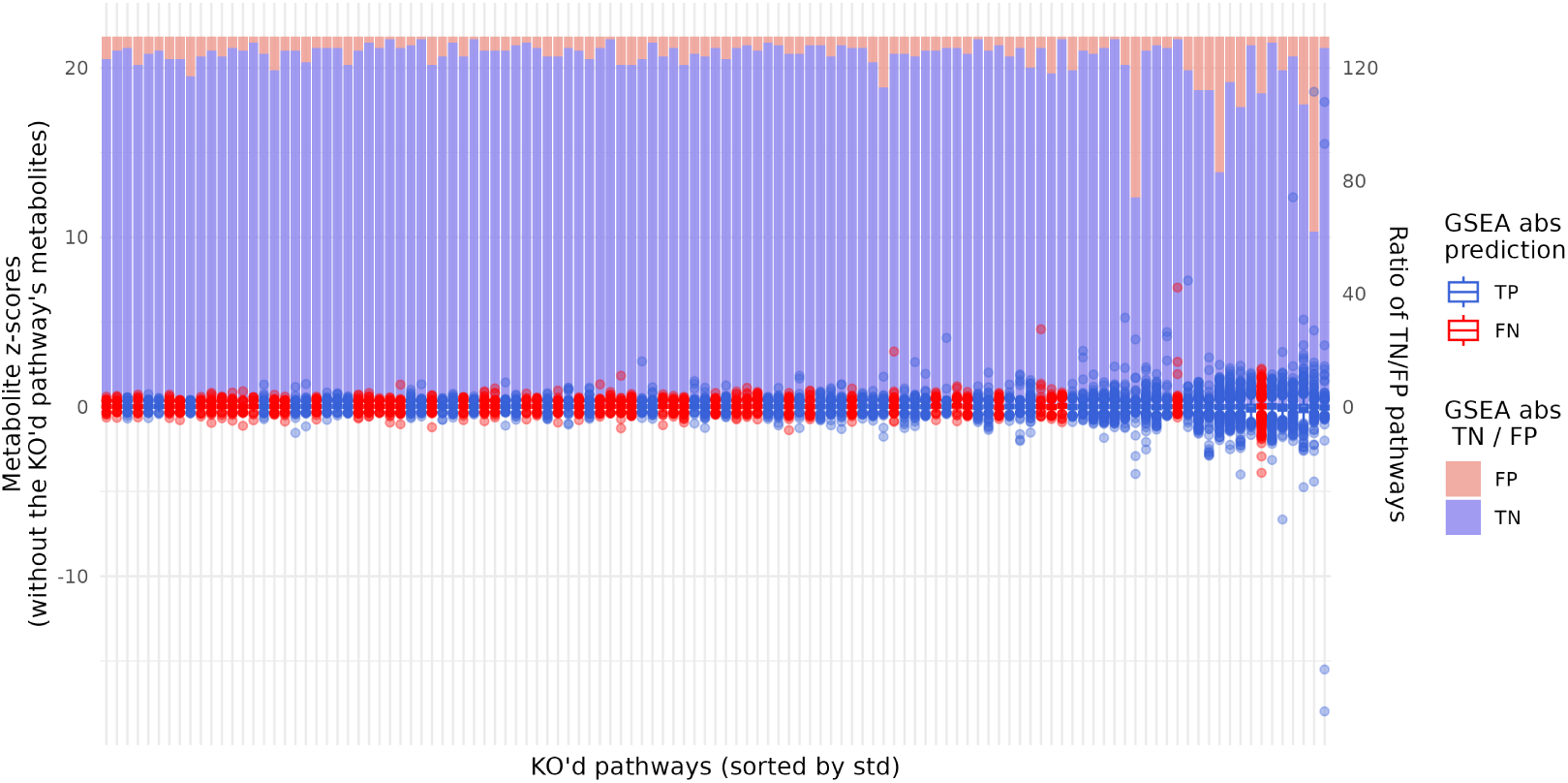
Boxplots of metabolite z-scores for each knocked out pathway. Pathways (x-axis) are sorted by standard deviation, and coloured by GSEA prediction (TP/FN). The y-axis shows the z-scores of all metabolites except those in the knocked out pathway. Outliers are shown as points. The secondary y-axis (right) shows the number of FP and TN for that pathway knockout.

**Fig E in S1 Text:**
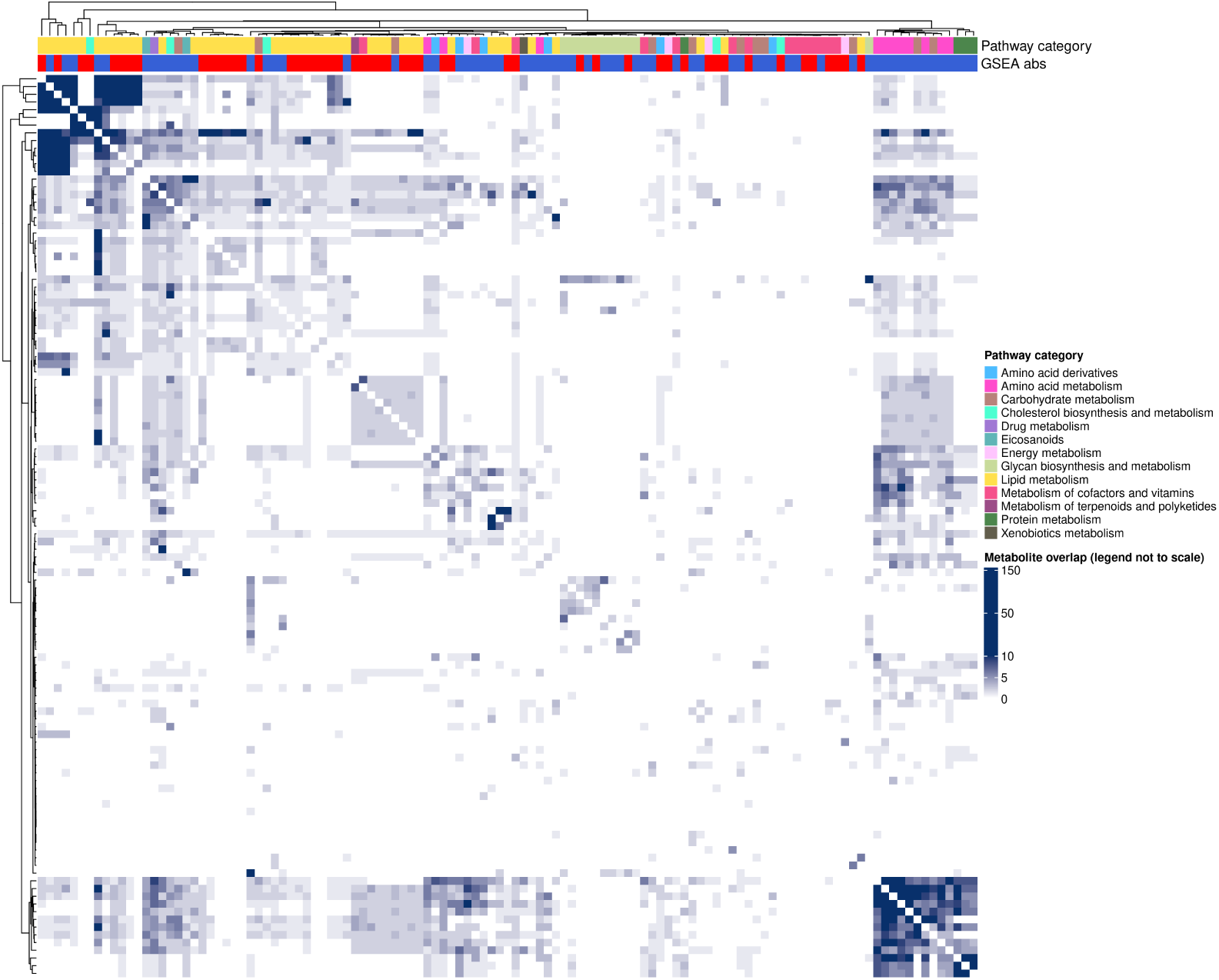
Overlap heatmap. Overlap between all pairs of pathways in the metabolic network (calculated when generating the PathwayNet graph). The idea is that the more a pathway shares metabolites with other pathways, the more likely it is to not be enriched when knocked out, due to compensating fluxes and less specific metabolic profiles. The darker cells representing more shared metabolites between the two pathways. The top left quarter pathways share many metabolites among them. The majority of these pathways are annotated as lipid pathways, many of which are FNs. The pathways in the bottom right cluster appear to group cofactor and vitamins pathways together, which are able to be predicted well.

**Fig F in S1 Text:**
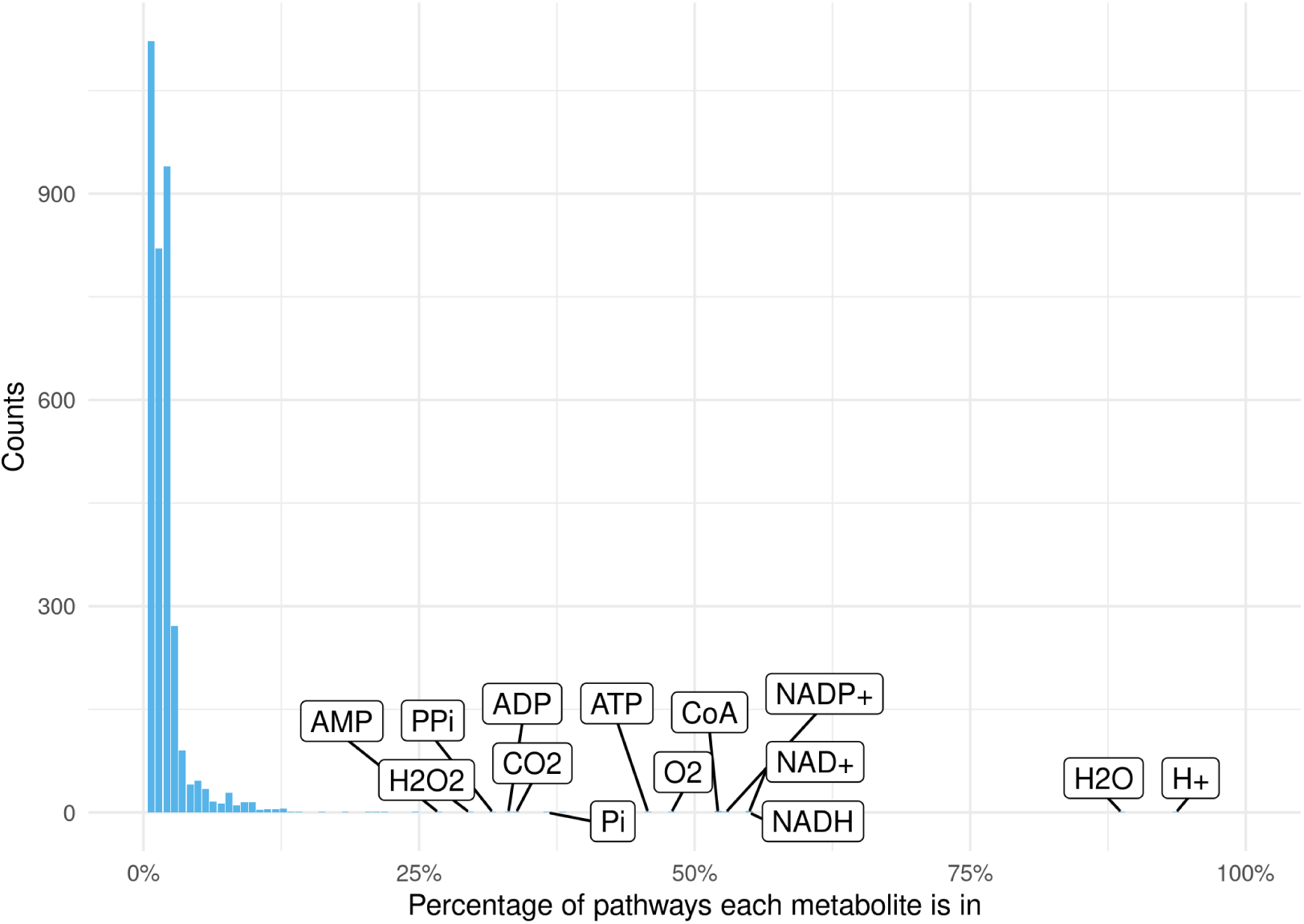
Distribution of the percentage of pathways each metabolite is in. Metabolites which are in many pathways are side compounds, such as H+, H2O, and NADH.

**Fig G in S1 Text:**
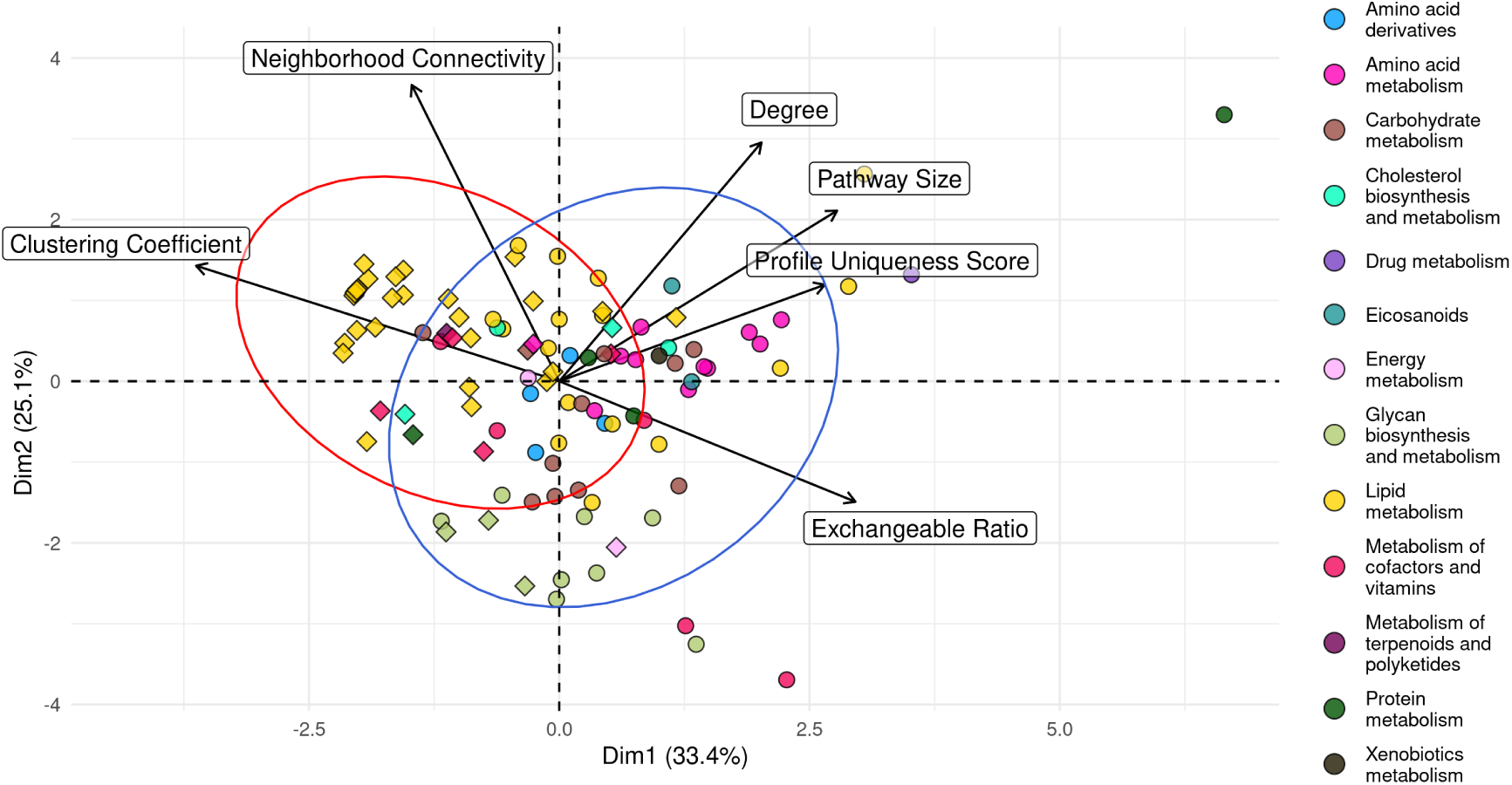
PCA of pathways using 6 metrics and properties. TP pathways are shown as circles, FN pathways are shown as diamonds. Pathway colour is the pathway category. When coloured by pathway category, similar clusters to the pathway graph appear. For example, Amino acid metabolism pathways tend to be large in size while also having a relatively high metabolic profile uniqueness score, and a low clustering coefficient. This reveals that pathways with the same super-category, and therefore somewhat similar functions, tend to have similar properties. Most of the FN pathways outside of the blue (TP) ellipse are lipid pathways, which shows that these are the pathways that are the most different in terms of pathway properties to the TP pathways.

#### Application to Recon2.2

The following figures are reproduced with Recon2.2 instead of Human1 as seen in the main text.

**Fig H in S1 Text:**
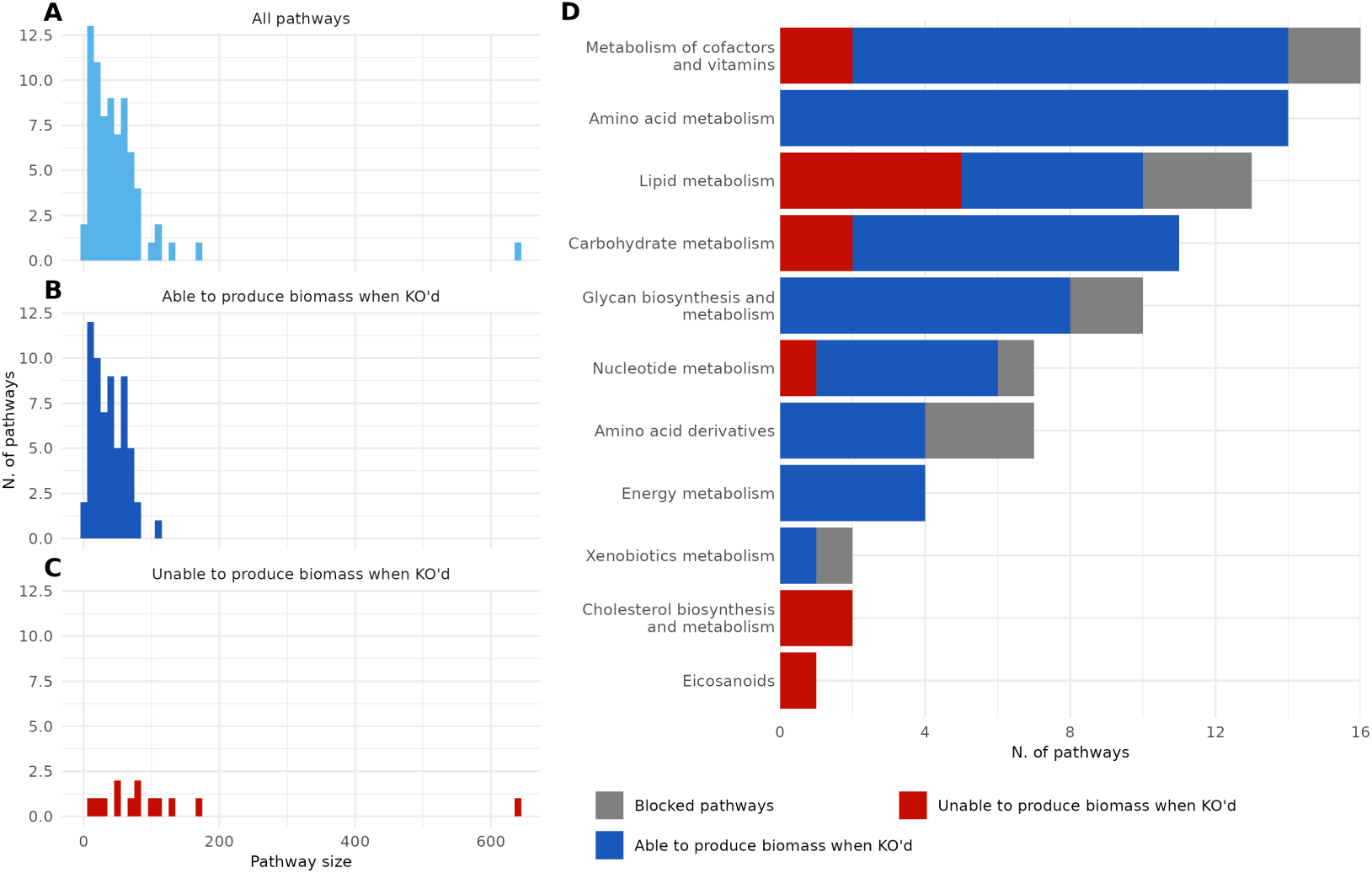
Fig 1, with Recon2.2 instead of Human1.

**Fig I in S1 Text:**
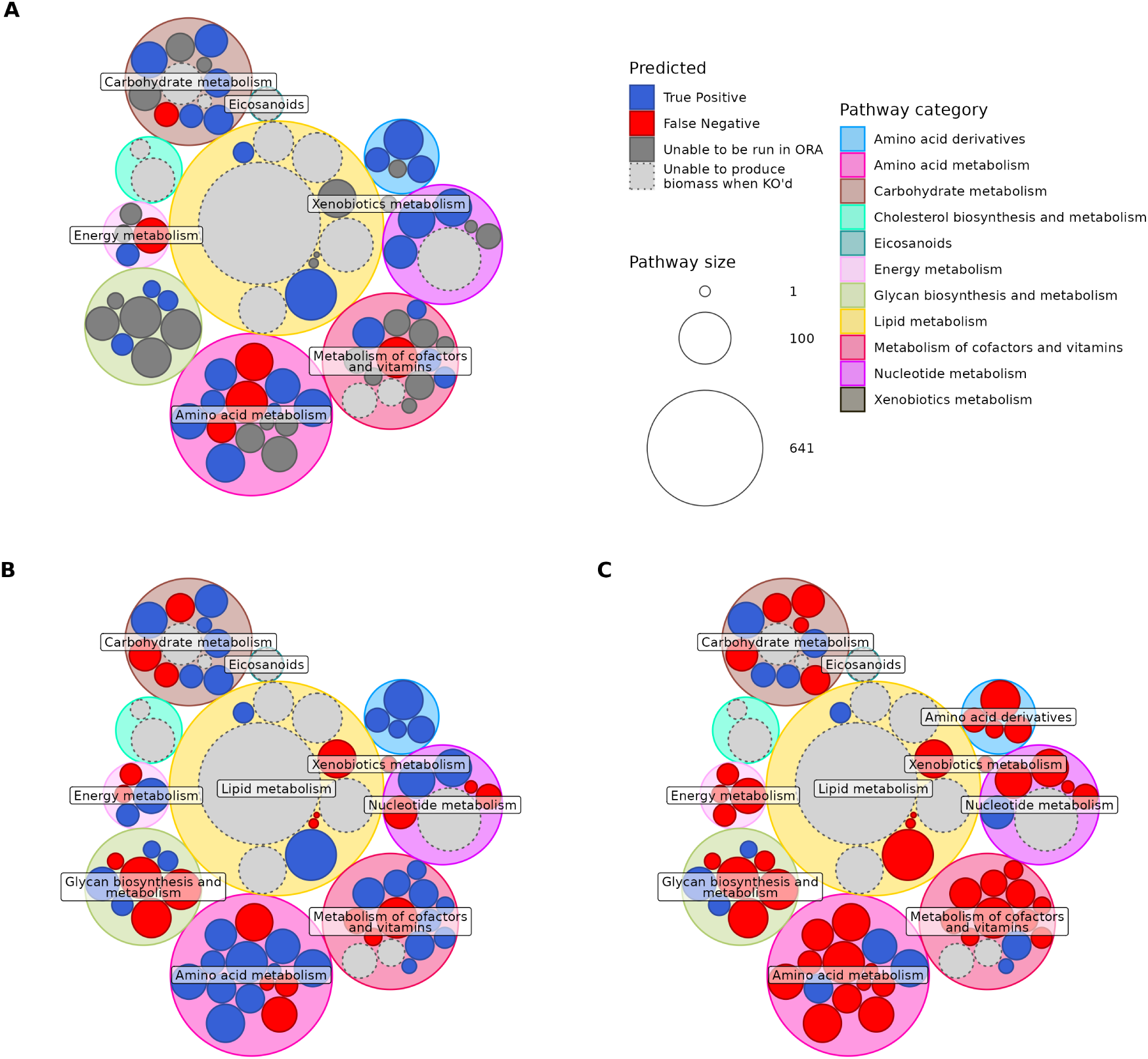
Fig 2, with Recon2.2 instead of Human1.

**Fig J in S1 Text:**
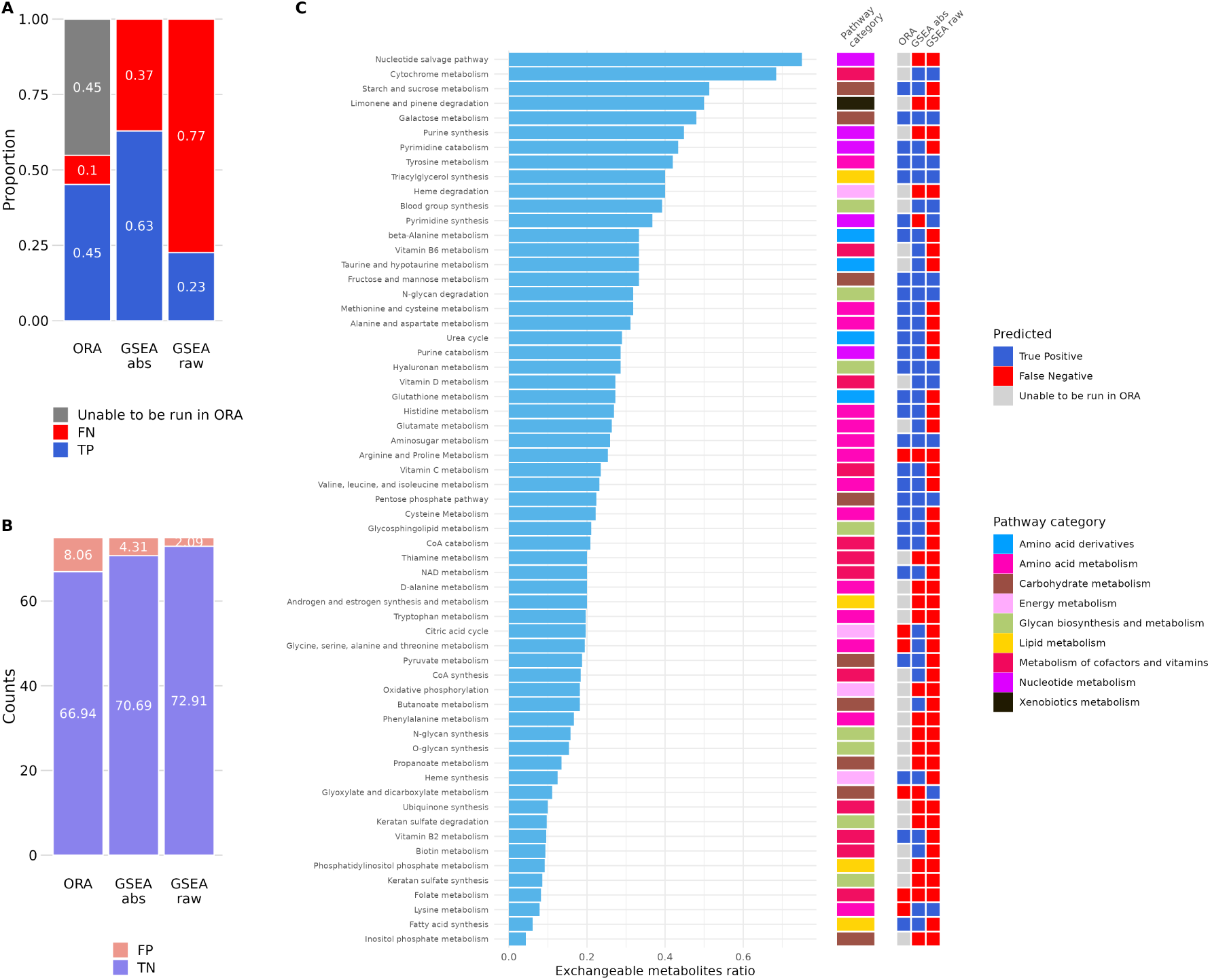
Fig 3 and 4, with Recon2.2 instead of Human1.

**Fig K in S1 Text:**
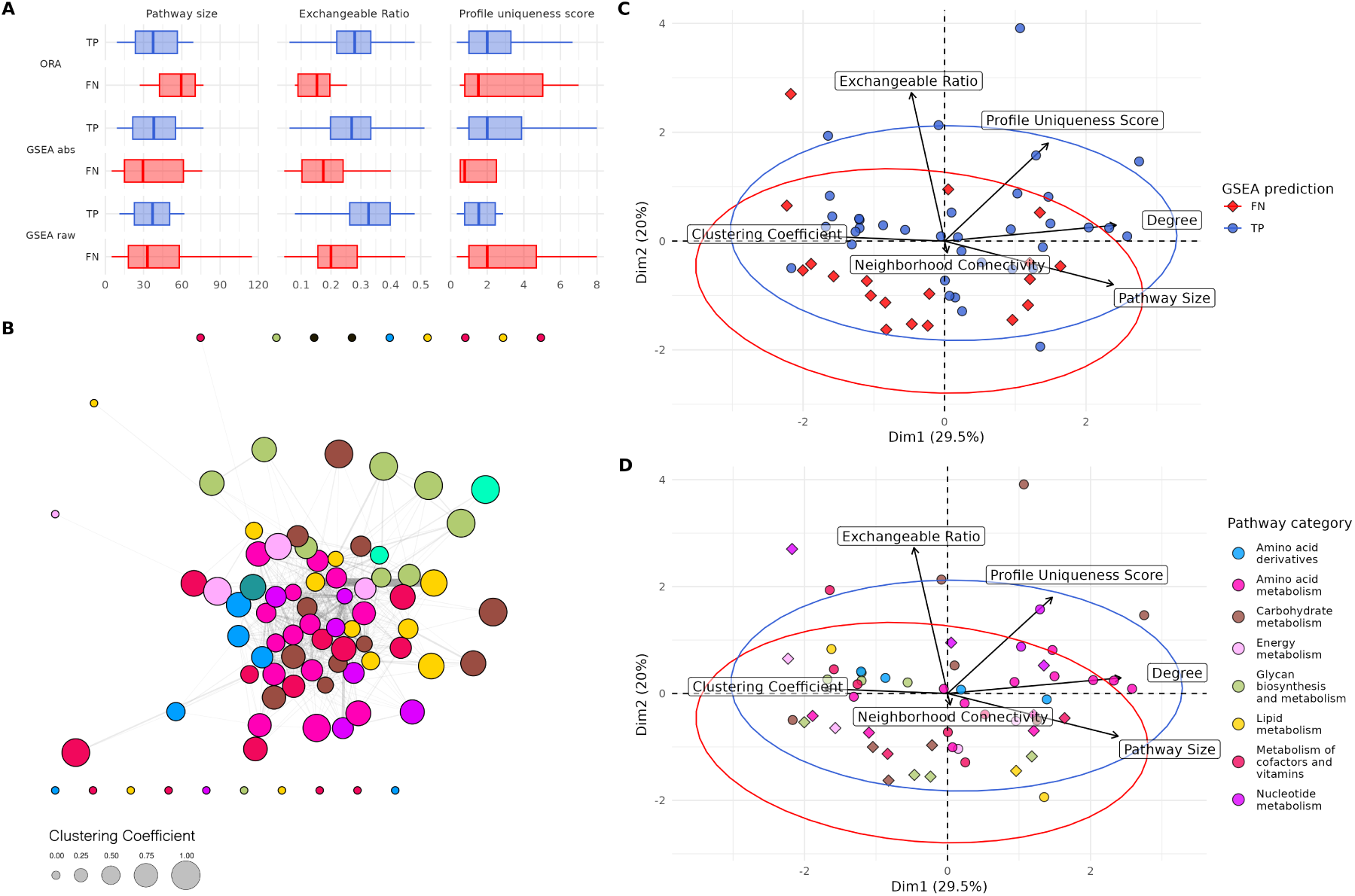
Fig 5, with Recon2.2 instead of Human1.

#### Application to KEGG

The following figures are reproduced with KEGG pathways instead of Human1 pathways as seen in the main text. The simulation results are from Human1 pathway KOs.

**Fig L in S1 Text:**
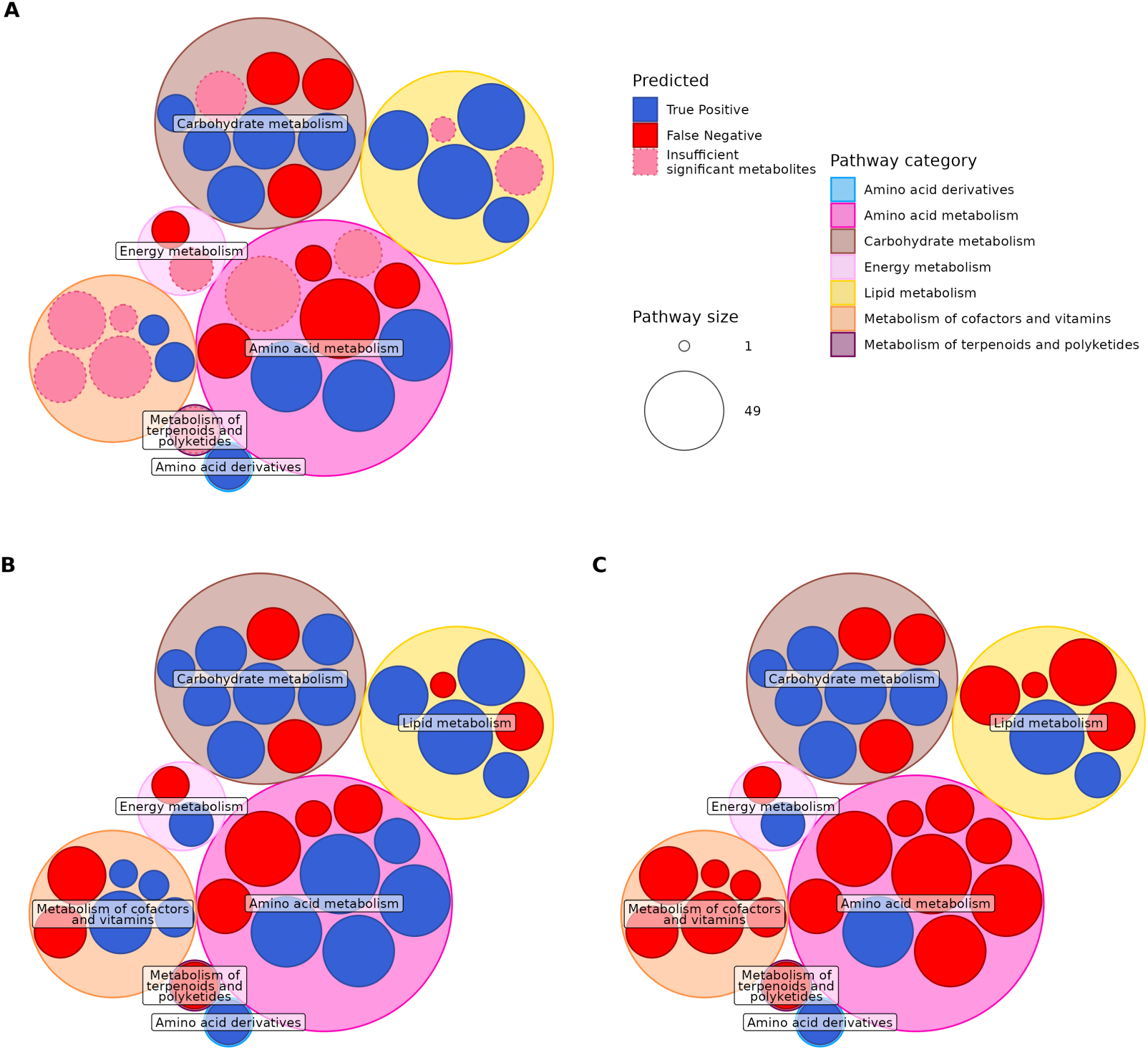
Fig 2, with KEGG pathways instead of Human1 pathways.

**Fig M in S1 Text:**
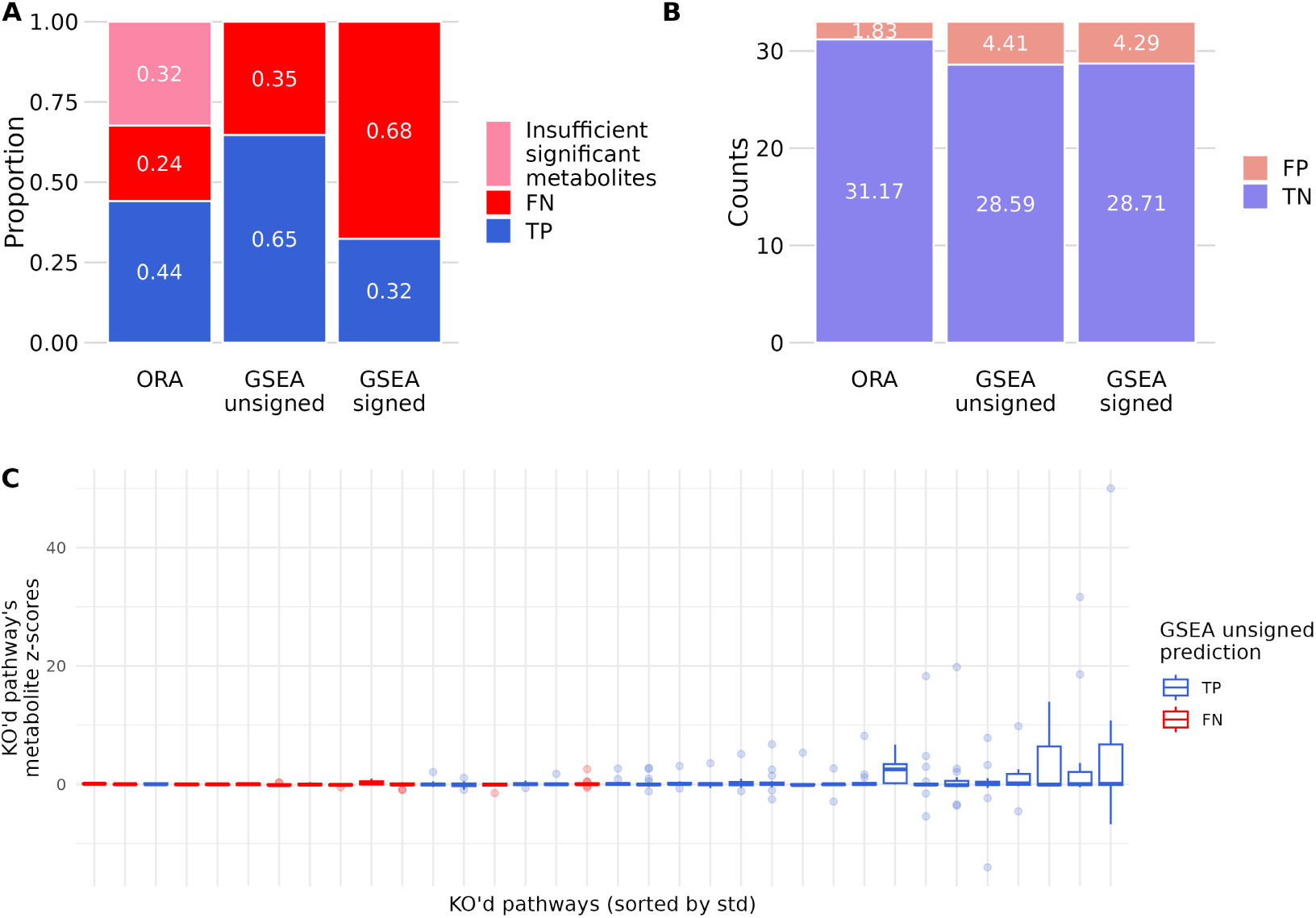
Fig 3, with KEGG pathways instead of Human1 pathways.

## References

Abatangelo, L., Maglietta, R., Distaso, A., D’Addabbo, A., Creanza, T.M., Mukherjee, S., Ancona, N. (2009, September). Comparative study of gene set enrichment methods. BMC Bioinformatics, 10 (1), 275, 10.1186/1471-2105-10-275 Retrieved 2024-09-20, from https://doi.org/10.1186/1471-2105-10-275

Chagoyen, M., & Pazos, F. (2011, March). MBRole: enrichment analysis of metabolomic data. Bioinformatics (Oxford, England), 27 (5), 730–731, 10.1093/ bioinformatics/btr001

Cooke, J., Delmas, M., Wieder, C., Mier, P.R., Frainay, C., Vinson, F., . . . Jourdan, F. (2024, February). Genome scale metabolic network modelling for metabolic profile predictions. PLOS Computational Biology, 20 (2), e1011381, 10.1371/journal.pcbi.1011381 Retrieved 2024-05-07, from https://journals.plos.org/ploscompbiol/article?id=10.1371/journal.pcbi.1011381 (Publisher: Public Library of Science)

Fang, Z., Liu, X., Peltz, G. (2023, January). GSEApy: a comprehensive package for performing gene set enrichment analysis in Python. Bioinformatics (Oxford, England), 39 (1), btac757, 10.1093/bioinformatics/btac757

Kanehisa, M., & Goto, S. (2000, January). KEGG: kyoto encyclopedia of genes and genomes. Nucleic Acids Research, 28 (1), 27–30, 10.1093/nar/28.1.27

Karnovsky, A., & Li, S. (2020). Pathway Analysis for Targeted and Untargeted Metabolomics. Methods in Molecular Biology (Clifton, N.J.), 2104, 387–400, 10.1007/978-1-0716-0239-319

Liu, Q., Dinu, I., Adewale, A.J., Potter, J.D., Yasui, Y. (2007, November). Comparative evaluation of gene-set analysis methods. BMC Bioinformatics, 8 (1), 431, 10.1186/1471-2105-8-431 Retrieved 2024-09-20, from https://doi.org/10.1186/1471-2105-8-431

Marco-Ramell, A., Palau-Rodriguez, M., Alay, A., Tulipani, S., Urpi-Sarda, M., Sanchez-Pla, A., Andres-Lacueva, C. (2018, January). Evaluation and comparison of bioinformatic tools for the enrichment analysis of metabolomics data. BMC Bioinformatics, 19, 1, 10.1186/s12859-017-2006-0 Retrieved 2024-08-13, from https://www.ncbi.nlm.nih.gov/pmc/articles/PMC5749025/

Nguyen, T.-M., Shafi, A., Nguyen, T., Draghici, S. (2019, October). Identifying significantly impacted pathways: a comprehensive review and assessment. Genome Biology, 20 (1), 203, 10.1186/s13059-019-1790-4 Retrieved 2024-09-20, from https://doi.org/10.1186/s13059-019-1790-4

Robinson, J.L., Kocabaş, P., Wang, H., Cholley, P.-E., Cook, D., Nilsson, A., . . . Nielsen, J. (2020, March). An atlas of human metabolism. Science Signaling, 13 (624), eaaz1482, 10.1126/scisignal.aaz1482

Shannon, P., Markiel, A., Ozier, O., Baliga, N.S., Wang, J.T., Ramage, D., . . . Ideker, T. (2003, November). Cytoscape: a software environment for integrated models of biomolecular interaction networks. Genome Research, 13 (11), 2498–2504, 10.1101/gr.1239303

Subramanian, A., Tamayo, P., Mootha, V.K., Mukherjee, S., Ebert, B.L., Gillette, M.A., . . . Mesirov, J.P. (2005, October). Gene set enrichment analysis: a knowledge-based approach for interpreting genome-wide expression profiles. Proceedings of the National Academy of Sciences of the United States of America, 102 (43), 15545–15550, 10.1073/pnas.0506580102

Swainston, N., Smallbone, K., Hefzi, H., Dobson, P.D., Brewer, J., Hanscho, M., . . . Mendes, P. (2016). Recon 2.2: from reconstruction to model of human metabolism. Metabolomics, 12, 109, 10.1007/s11306-016-1051-4 Retrieved 2023-10-18, from https://www.ncbi.nlm.nih.gov/pmc/articles/PMC4896983/

Thiele, I., Swainston, N., Fleming, R.M.T., Hoppe, A., Sahoo, S., Aurich, M.K., . . . Palsson, B.O. (2013, May). A community-driven global reconstruction of human metabolism. Nature Biotechnology, 31 (5), 419–425, 10.1038/nbt.2488 Retrieved 2022-07-25, from https://www.nature.com/articles/nbt.2488 Number: 5 Publisher: Nature Publishing Group)

Wieder, C., Frainay, C., Poupin, N., Rodŕıguez-Mier, P., Vinson, F., Cooke, J., . . . Ebbels, T. (2021, September). Pathway analysis in metabolomics: Recommendations for the use of over-representation analysis. PLOS Computational Biology, 17 (9), e1009105, 10.1371/journal.pcbi.1009105 Retrieved 2024-01-17, from https://journals.plos.org/ploscompbiol/article?id=10.1371/journal.pcbi.1009105 (Publisher: Public Library of Science)

Wieder, C., Lai, R.P.J., Ebbels, T.M.D. (2022, November). Single sample pathway analysis in metabolomics: performance evaluation and application. BMC Bioinformatics, 23 (1), 481, 10.1186/s12859-022-05005-1 Retrieved 2024-08-05, from https://doi.org/10.1186/s12859-022-05005-1

Yu, C., Woo, H.J., Yu, X., Oyama, T., Wallqvist, A., Reifman, J. (2017, October). A strategy for evaluating pathway analysis methods. BMC Bioinformatics, 18 (1), 453, 10.1186/s12859-017-1866-7 Retrieved 2024-09-20, from https://doi.org/10.1186/s12859-017-1866-7

