## Supplementary Information for "Simulated metabolic profiles reveal biases in pathway analysis methods"

### S1 Text

#### ***Methods in S1 Text***

The *clustering coefficient* quantifies how connected each pathway's neighbouring pathways are. For a node, it is defined as the actual number of links between its neighbouring nodes divided by the maximum possible number of links that could exist between these nodes.

For a pathway $p_i$:

$$p_i\text{'s clustering coefficient} = \frac{2 * \text{n. edges in } N_i}{k_i(k_i - 1)} \tag{1}$$

where $N_i$ is $p_i$'s (immediate) neighbourhood, and $k_i$ is the number of nodes in $N_i$.

The *neighbourhood connectivity* is defined as the average connectivity of all of a node's neighbours. A node's connectivity, or degree, is the number of connected edges, in our case the number of neighbours.

$$p_i\text{'s neighbourhood connectivity} = \frac{\sum_{n=1}^{k_i} \deg(N_{i,n})}{k_i} \tag{2}$$

where $N_i$ is $p_i$'s (immediate) neighbourhood, $N_{i,n}$ is the $n$th node in $N_i$, and $k_i$ is the number of nodes in $N_i$.

#### ***Eq A in S1 Text***

$$\begin{gathered}
45\text{ ATP } + 0.0267\text{ DNA } + 45\text{ H2O } + 0.1124\text{ RNA} \\
+0.0012\text{ cofactor pool biomass } + 0.4062\text{ glycogen } + 0.2212\text{ lipid pool biomass} \\
+0.4835\text{ metabolite pool biomass } + 5.3375\text{ protein pool biomass} \\
\downarrow \\
45\text{ ADP } + 45\text{ H+ } + 45\text{ Pi } + \text{ biomass}
\end{gathered} \tag{3}$$

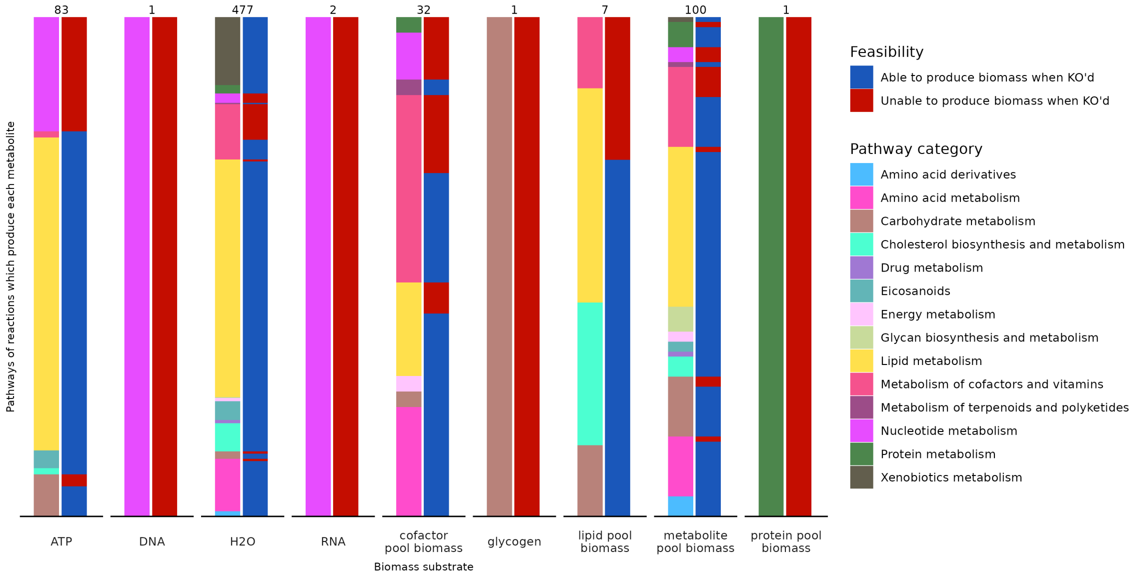


**Fig A in S1 Text**: Pathway categories of reactions which produce each substrate in the biomass reaction. Each bar is the total number of reactions that produce that metabolite prior to the bioomass reaction. The left bars are the pathway super-categories for those reactions, and the right bars are whether those pathways are infeasible (red) or functional (blue).

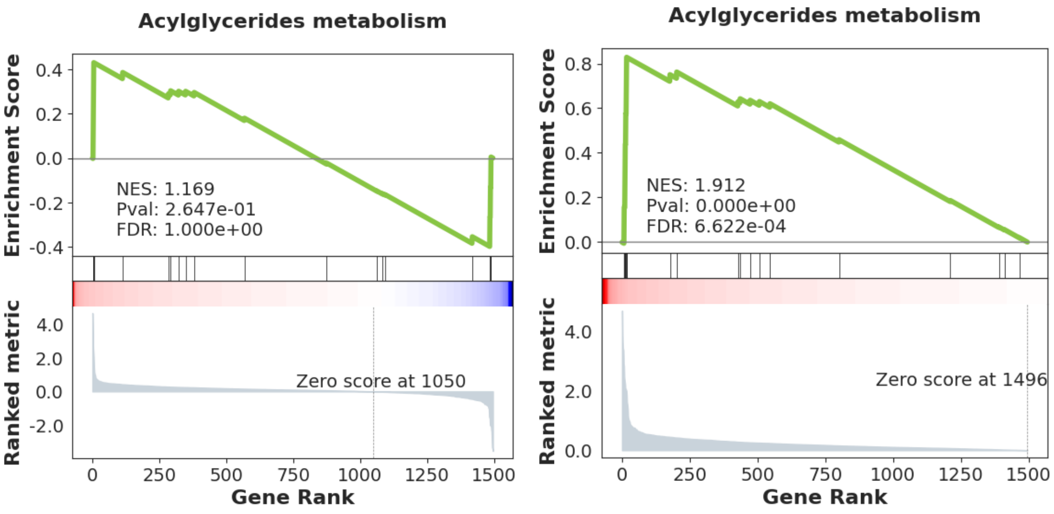


(a) GSEA with signed z-scores.

(b) GSEA with unsigned (absolute) z-scores.

**Fig B in S1 Text**: GSEA enrichment plots for Acylglycerides metabolism with GSEA abs and GSEA raw. The signed z-scores result in a split profile (similar contributions of high and low abundance metabolites), and a non-significant enrichment. By converting the z-scores to absolute values, they are clustered together in the ranking list, resulting in a correct significant enrichment.

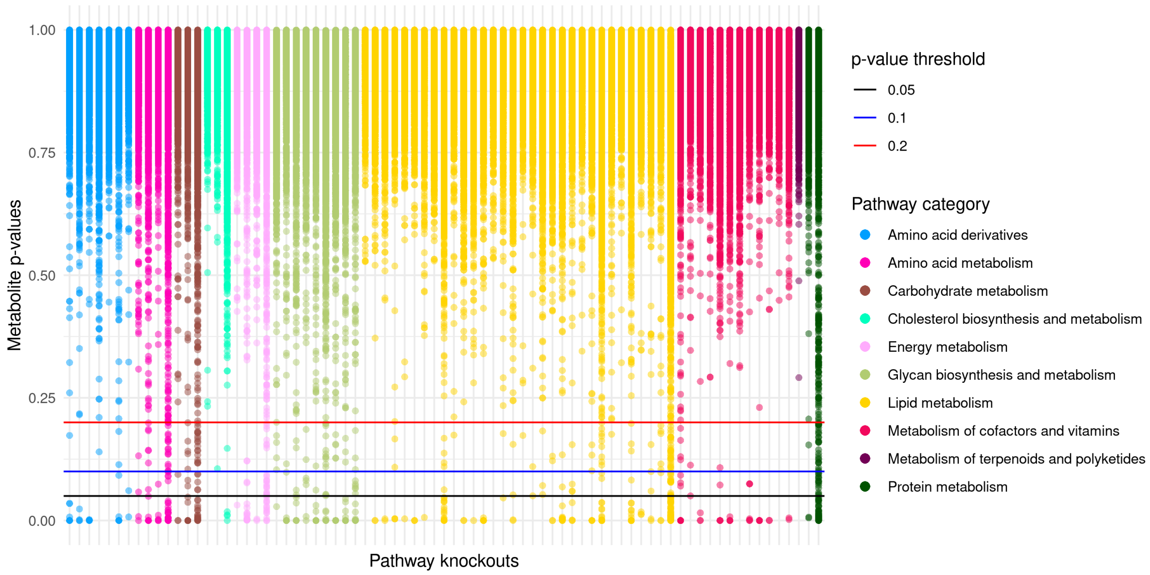


**Fig C in S1 Text**: Different DA (differentially abundant) thresholds on all pathway z-scores for ORA. We used $p \leq 0.05$ in our analyses. Pathways are grouped by super category.

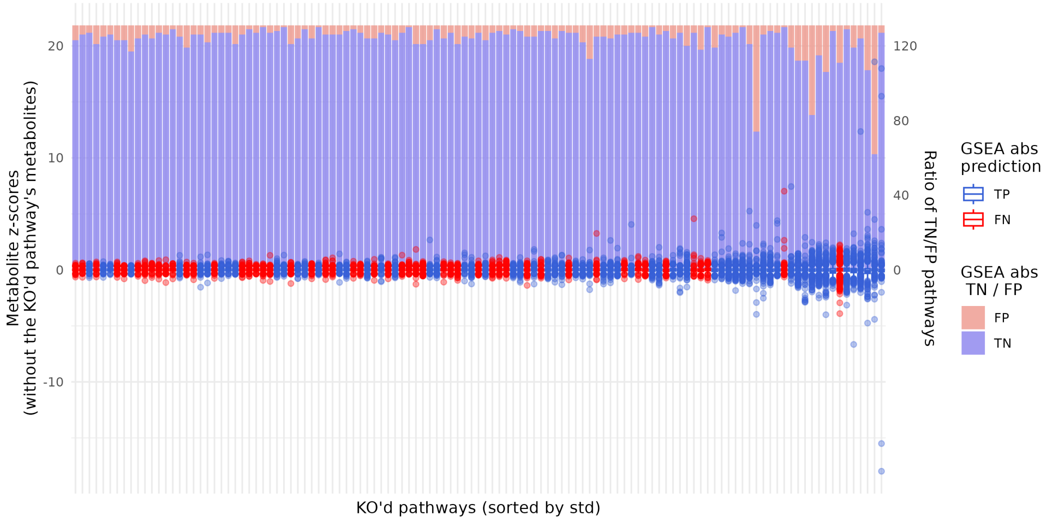


**Fig D in S1 Text**: Boxplots of metabolite z-scores for each knocked out pathway. Pathways (x-axis) are sorted by standard deviation, and coloured by GSEA prediction (TP/FN). The y-axis shows the z-scores of all metabolites except those in the knocked out pathway. Outliers are shown as points. The secondary y-axis (right) shows the number of FP and TN for that pathway knockout.

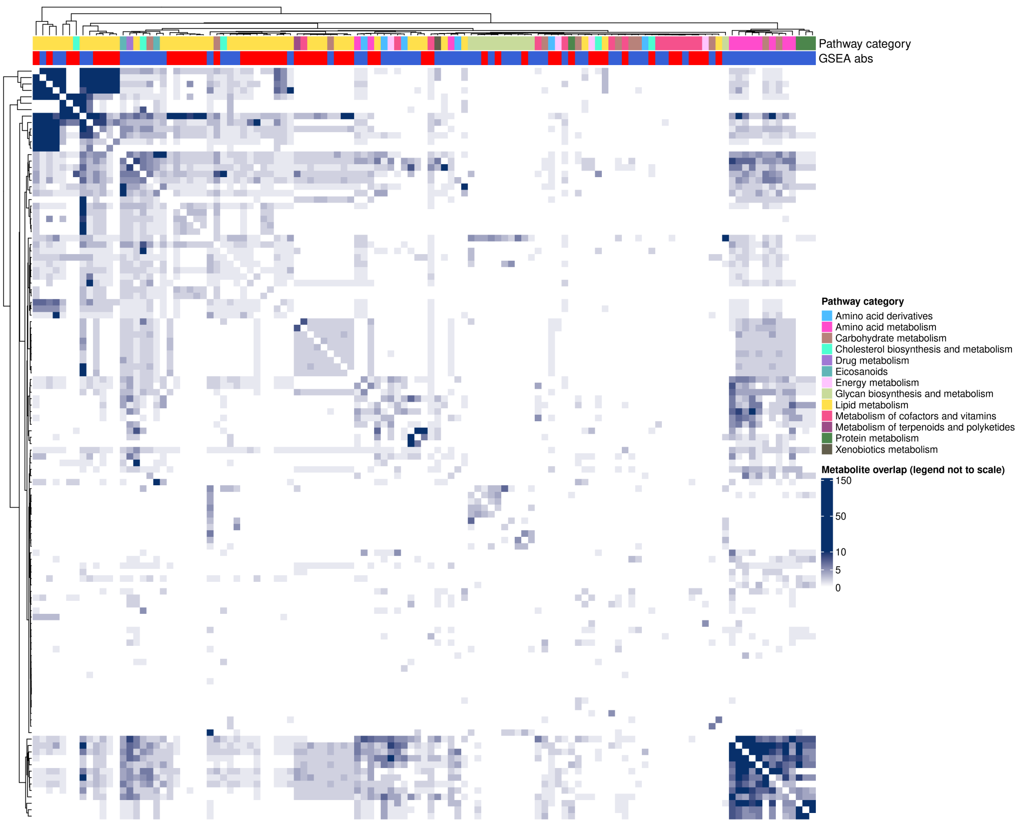


**Fig E in S1 Text**: Overlap heatmap. Overlap between all pairs of pathways in the metabolic network (calculated when generating the PathwayNet graph). The idea is that the more a pathway shares metabolites with other pathways, the more likely it is to not be enriched when knocked out, due to compensating fluxes and less specific metabolic profiles. The darker cells representing more shared metabolites between the two pathways. The top left quarter pathways share many metabolites among them. The majority of these pathways are annotated as lipid pathways, many of which are FNs. The pathways in the bottom right cluster appear to group cofactor and vitamins pathways together, which are able to be predicted well.

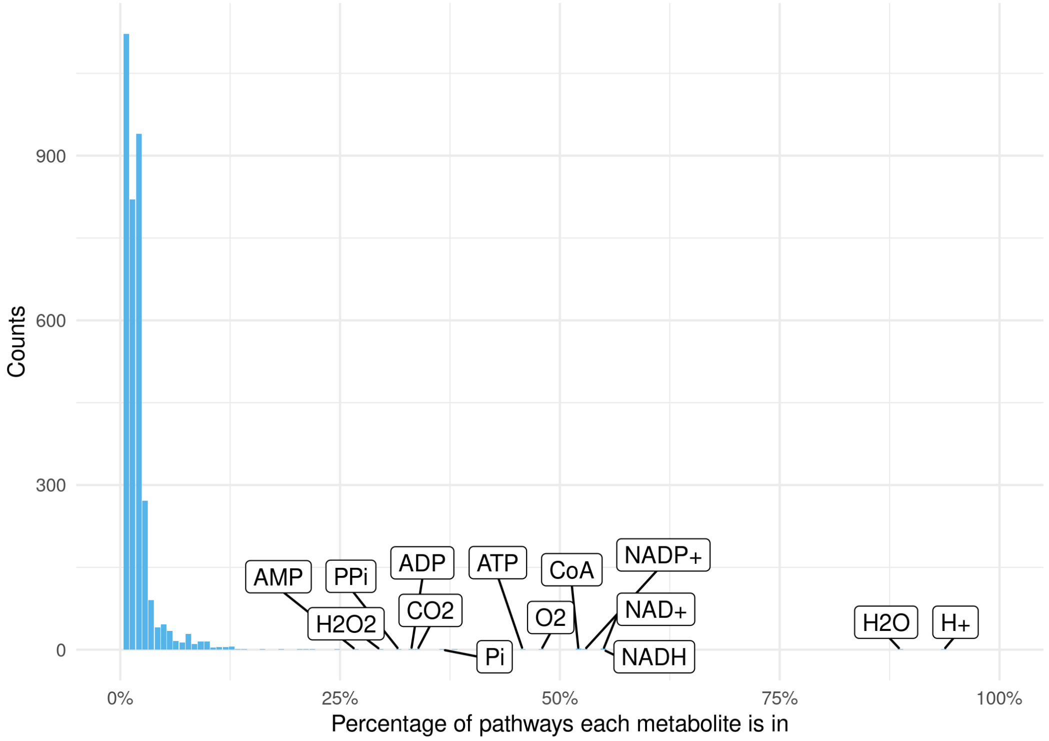


**Fig F in S1 Text**: Distribution of the percentage of pathways each metabolite is in. Metabolites which are in many pathways are side compounds, such as H+, H2O, and NADH.

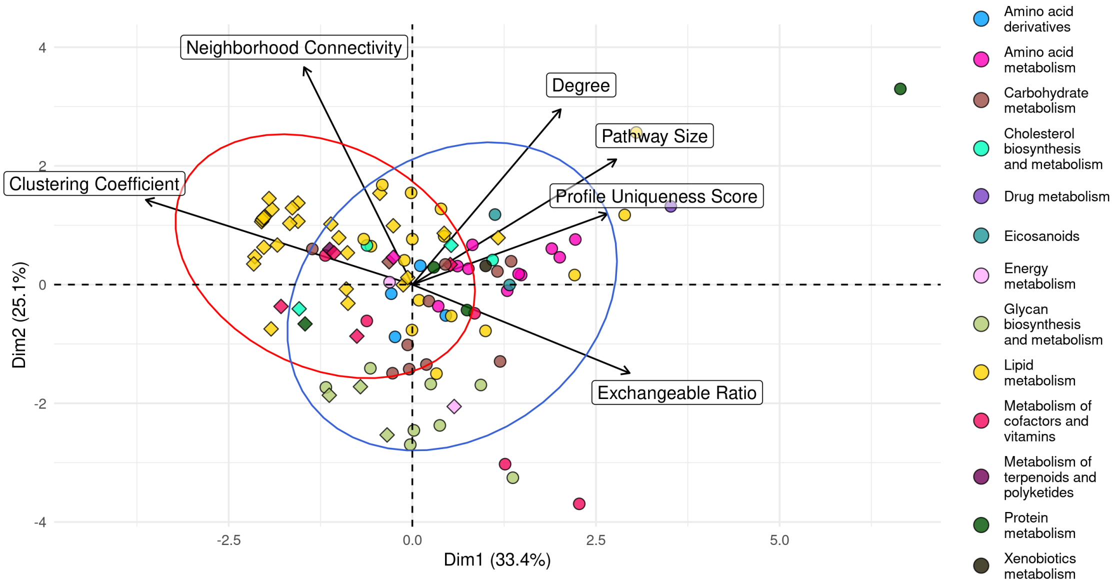


**Fig G in S1 Text**: PCA of pathways using 6 metrics and properties. TP pathways are shown as circles, FN pathways are shown as diamonds. Pathway colour is the pathway category. When coloured by pathway category, similar clusters to the pathway graph appear. For example, Amino acid metabolism pathways tend to be large in size while also having a relatively high metabolic profile uniqueness score, and a low clustering coefficient. This reveals that pathways with the same super-category, and therefore somewhat similar functions, tend to have similar properties. Most of the FN pathways outside of the blue (TP) ellipse are lipid pathways, which shows that these are the pathways that are the most different in terms of pathway properties to the TP pathways.

#### *Application to Recon2.2*

The following figures are reproduced with Recon2.2 instead of Human1 as seen in the main text.

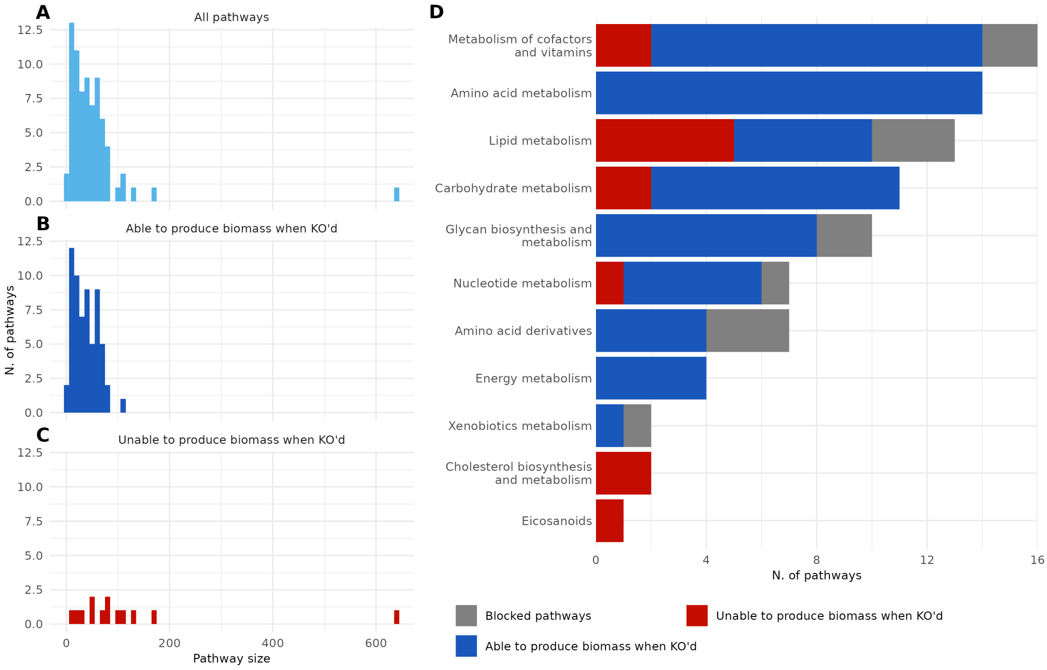


**Fig H in S1 Text**: Fig 1, with Recon2.2 instead of Human1.

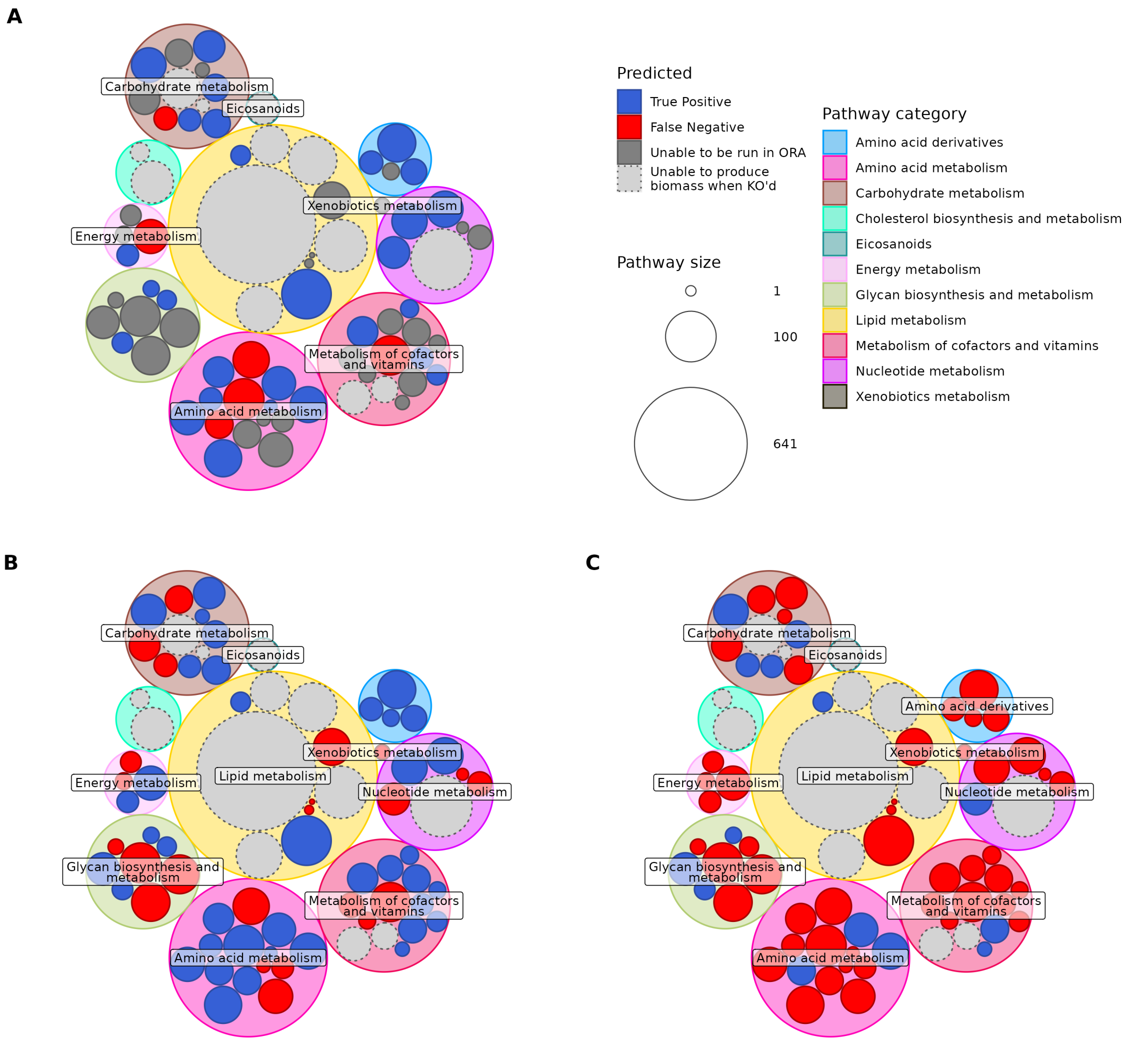


**Fig I in S1 Text**: Fig 2, with Recon2.2 instead of Human1.

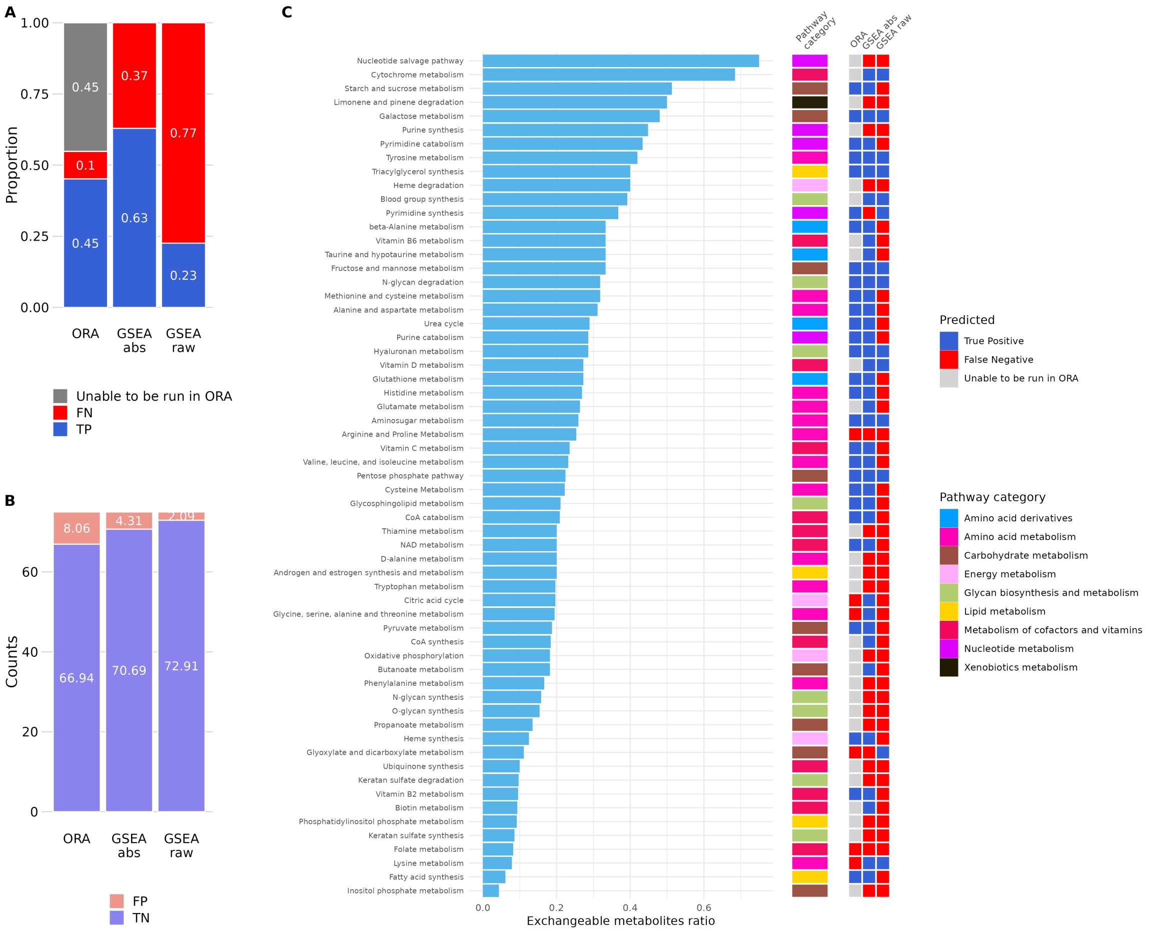


**Fig J in S1 Text**: Fig 3 and 4, with Recon2.2 instead of Human1.

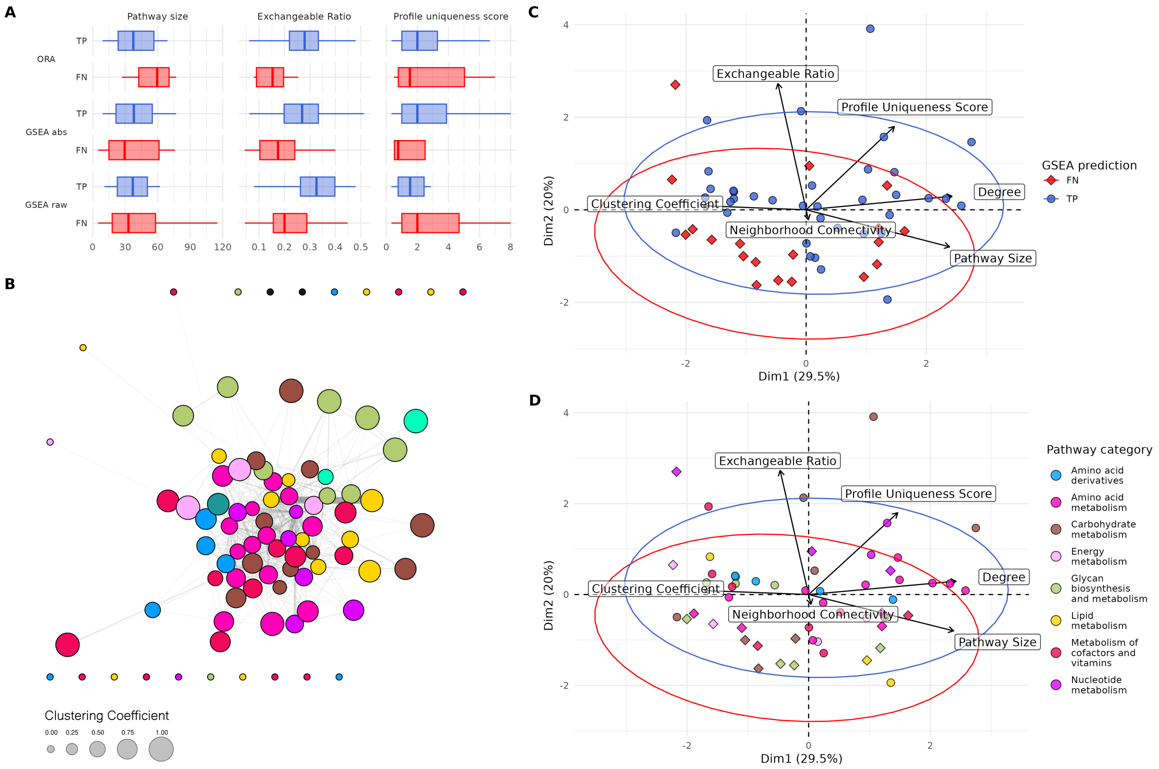


**Fig K in S1 Text**: Fig 5, with Recon2.2 instead of Human1.

#### *Application to KEGG*

The following figures are reproduced with KEGG pathways instead of Human1 pathways as seen in the main text. The simulation results are from Human1 pathway KOs.

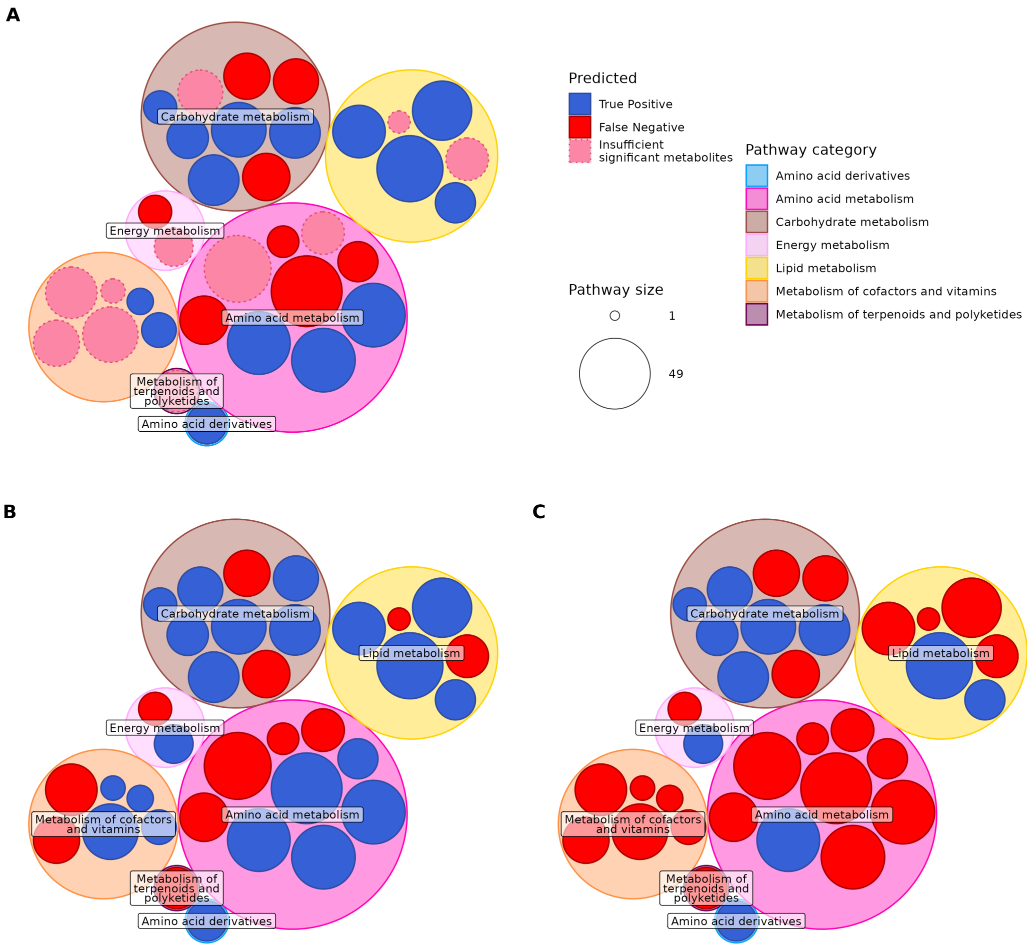


**Fig L in S1 Text**: Fig 2, with KEGG pathways instead of Human1 pathways.

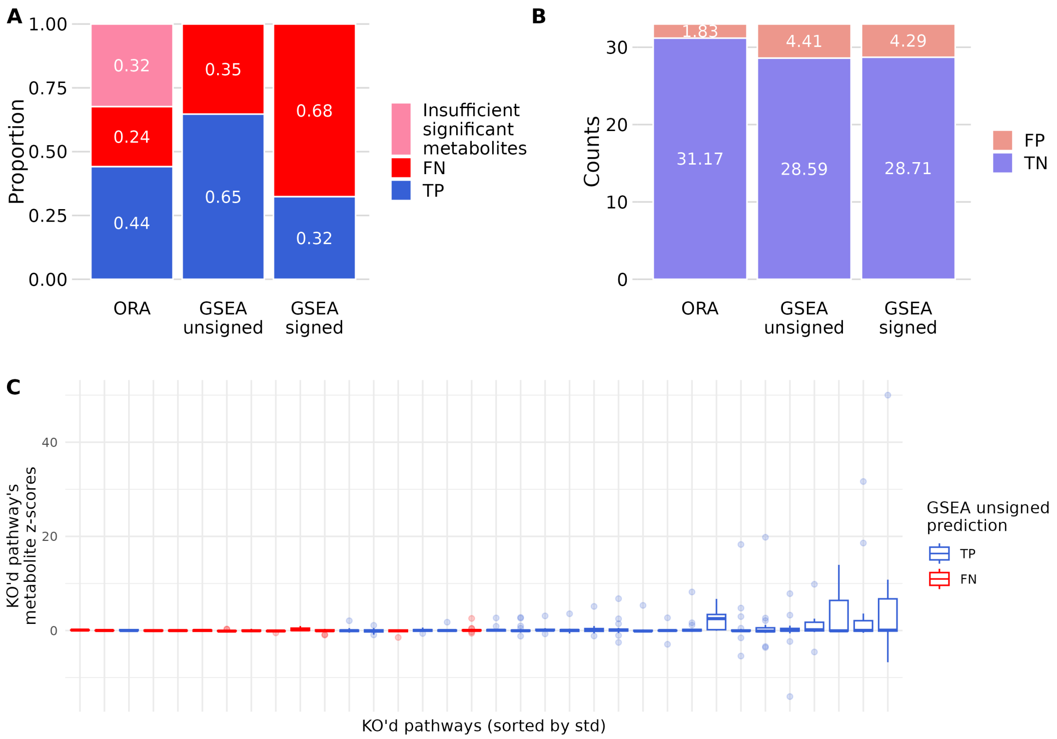


**Fig M in S1 Text**: Fig 3, with KEGG pathways instead of Human1 pathways.
